# Markers, Mechanisms and Metrics of Biological Aging: A Scoping Review

**DOI:** 10.1101/2024.10.29.620898

**Authors:** Alison Ziesel, Jennifer Reeves, Anastasia Mallidou, Lorelei Newton, Ryan Rhodes, Jie Zhang, Theone Paterson, Hosna Jabbari

**Affiliations:** Department of Biomedical Engineering, University of Alberta, Edmonton, Alberta, Canada; Department of Psychology, University of Victoria, Victoria, BC, Canada; School of Nursing, University of Victoria, Victoria, BC, Canada; Department of Exercise Science, Physical & Health Education, University of Victoria, Victoria, BC, Canada; Gustavson School of Business, University of Victoria, Victoria, BC, Canada

## Abstract

Biological aging is a rapidly growing area of research, which entails characterizing the rate of aging independent of an individual’s chronological age. In this review, we analyze the results of biological aging research in 435 papers published in a twelve year window, revealing changing patterns of molecular markers of biological aging use over time, and the development of novel metrics of biological aging. We further identify consistent and discordant research findings, as well as areas of potential future research focusing on questions of measurement with methylation or biomarker-based assessment and other variables relevant to the study of biological age.

## Introduction

Biological aging is the process by which organisms develop, mature, and age, in parallel with but not exclusively dependent upon their chronological age. As biological aging is distinct from chronological age, landmarks of development and onset of age-related diseases may not be accurately predicted based on chronological age alone. Moreover, the further an organism progresses from birth, the broader the difference between chronological and biological age may become, meaning that later in life the biological age differential may be quite significant. For these reasons it is important to be able to assess the progress of biological aging to more precisely determine these intervals of developmental change and disease vulnerability.

Biological aging is characterized by changes from the subcellular level to the organismal level. Such diverse features as telomere attrition and hair whitening have been positively correlated with advancing age, and so the process of biological aging may be observed and measured both microscopically and individually. These measurable features may be indicative of an individual entering a stage of life relatively advanced for their chronological age, in which they are prematurely more susceptible to a variety of age-related maladies. Being able to assess individuals for their biological age and changes in their rates of aging may allow for intervention to take place before disease onset, making accurate measurement of biological age critical for identifying windows for therapeutic and preventative measures to ensure or promote continuing good health.

The questions of what biological age represents, and how biological aging occurs, have been investigated for decades. Initially characterized as ’functional age’ in studies conducted during the 1950s [273], biological age resurged as a serious inquiry in the 1960s and 1970s with the work of Dirken and Furukawa [93, 121], although there was some debate as to the utility of biological age as a marker of organismal development in the following decade [73, 87]. Biomarkers in the form of clinical parameters, or blood- or tissue-derived protein or metabolite levels were often employed in both these early studies and their use persisted over time. The early 1990s heralded the observation that telomeres in normal somatic cells lacking telomerase decrease with progressive cell cycles, suggesting that telomere length may be a usable proxy for cellular aging rates [224]. In the following decade it was discovered that mitochondrial DNA (mtDNA) levels decline with age in skeletal muscle, and later in other tissues including pancreatic islet cells [76, 434]. While initially reported as early as 1967, both global and site-specific methylation changes with respect to age were revisited in the early twenty first century, leading to a greater focus on these epigenetic changes in subsequent studies [33, 39, 112, 341]. This led to the development of the first epigenetic clocks in the 2010s, which undertake to predict age solely from the methylation status of a subset of cytosine-phosphate-guanosine (CpG) sites in the genome [147, 166]. These developments were followed by second generation epigenetic clocks, many of which incorporate additional, non-methylation data to predict biological age and in some cases, mortality [223, 245].

The preceding represents an exceedingly high-level overview of decades of work and as such is insufficient to adequately describe the wealth of discoveries and novel strategies developed over those years. Moreover, while there exist reviews that examine detailed subsets of this work, we did not identify a comprehensive review considering all molecular biological features of biological aging, along with the algorithms and clocks developed to interpret those data.

Key questions surrounding the measurement of biological age with respect to chronological age, or other features such as mortality, emerge throughout the body of literature assessed for this review. Some measurement questions are specific to certain methodologies, for example, methylation-based epigenetic changes have different considerations than do non-methylation molecular measures of aging. Other questions pertain to variables other than genetic and epigenetic: how do lifestyle choices and limitations modulate the course of healthy or pathologic aging in an individual?

To consider these questions, we undertook a scoping review study of those topics, focusing on works published during a twelve year window between 2011 and 2023. The strategy of a scoping review, rather than a systematic review, was employed for its capacity to expose relative strengths and shortcomings within a broader field of research. The identification of key questions and subtopics to pursue within the larger domain of molecular biological indicators, algorithmic approaches and their relationships and applicability to biological age determination is prioritized. Due to the breadth of research in the period analyzed, we have elected to portion our findings into a pair of review articles: this review detailing the molecular biological and algorithmic work on biological age measurement, with an additional review describing findings pertaining to clinical, socioeconomic and demographic features associated with changes in biological age.

## Materials and Methods

This review adhered to guidelines in the PRISMA Extension for Scoping Reviews [401], and was guided by Arksey and O’Malley’s methodological framework [9].

### Search Strategy

Four databases were searched from January 2011 to June 2023: PsycINFO, CINAHL, PubMed, and SPORTDiscus. Our search terms included on terms associated with biological aging and chronological aging, with all terms searched in the abstract and title. In addition, searches included relevant MeSH headings. Limits were applied to our search findings, specifically publication date (January 2011 to end of June 2023), subject (humans), age (adults aged 18 or older), and publication language (English).

### Eligibility Criteria

#### Population

To be eligible for inclusion, studies needed to include only human participants, aged 18 or older. These studies were included if they examined healthy adults or adults with specific health conditions (e.g., cancer). Studies including use of human-derived cell lines and human remains were retained. Studies which included non-human organisms or individuals under 18 years of age were excluded.

#### Outcomes

Studies chosen for inclusion measured chronological age and compare either a calculated biological age or a molecular indicator of biological aging (e.g., telomere length) with chronological age. Studies which examined the biological aging of only a specific organ were excluded.

#### Design

Empirical primary research studies were considered for inclusion if they were peer-reviewed and published in a journal. Non-research (e.g., opinion pieces, commentaries, letters, editorials), grey literature (e.g., theses, dissertations), theoretical papers, seminar papers, case studies, book chapters, study protocols, review articles, and articles in-press during the period assessed were excluded.

#### Other Considerations

In order to examine the most recent scientific findings, only studies published within a twelve and a half year window (January 2011 to the end of June 2023) were included. While only studies published in English were considered, studies were eligible for inclusion regardless of the country they were conducted in.

#### Study Selection and Data Charting

Reviewers used the software platform Covidence [75] to screen articles based on title and abstract. Two reviewers examined each title and abstract, and any conflicts during this step were resolved by a third reviewer. Full text articles were next screened according to the above described inclusion criteria. Conflicts were resolved by the three reviewers re-examining and discussing the article in question in the context of our inclusion criteria.

Data was extracted by reviewers, initially including the aims, population, methodology, and findings of the studies. Extracted data were then verified by one additional reviewer.

#### Data Synthesis

Following data extraction, a composite list of all outcomes which were compared to chronological and/or biological indicators of aging was composed. Subsequently, categories of those indicators employed were created; methodologies groups were created and adjusted as appropriate throughout the process. This resulted in 6 groups, which are described in Table 1. Due to the breadth of information summarized by our work, we have prepared two companion reviews: this review, which focuses on the underlying molecular biological features of biological aging, as well as the extant algorithms and clocks established for determining biological age, and an additional review, which focuses on the socioeconomic, physiological, and demographic parameters associated with changes in the biological aging rate.

**Table 1.** Categories and sub-categories of biological aging data analyzed in this review.

| Categories | Sub-categories |
| --- | --- |
| Molecular Biological Features | Mitochondrial DNA |
|  | Gene Expression |
|  | Non-Clock/Whole Genome Methylation |
|  | Telomere Length |
|  | Other Molecular Features |
| Biological Aging Algorithms and Clocks | Horvath’s Pan-Tissue Clock |
|  | Horvath’s Skin/Blood Clock |
|  | Hannum’s Clock |
|  | PhenoAge |
|  | GrimAge |
|  | Weidner’s Clock |
|  | Pace of Aging and DunedinPoAm |
|  | Klemera-Doubal Algorithm |
|  | Novel and Specific Clocks |

In our initial data collection, we identified 1244 works from the CINAHL, PsycINFO, PubMed and SPORTDiscus databases, from which 317 duplicates were removed (Figure 1). The remaining 927 records were screened for title and abstract, and 232 were removed. We sought 695 full text articles, of which one could not be retrieved, and excluded another 248 based on our inclusion and exclusion criteria. This resulted in 446 articles for our analyses. Prior to preparing this review, we excluded a further eleven papers as using exclusively non-molecular or non-biological metrics of biological aging, resulting in 435 papers analyzed.

**Fig 1.**
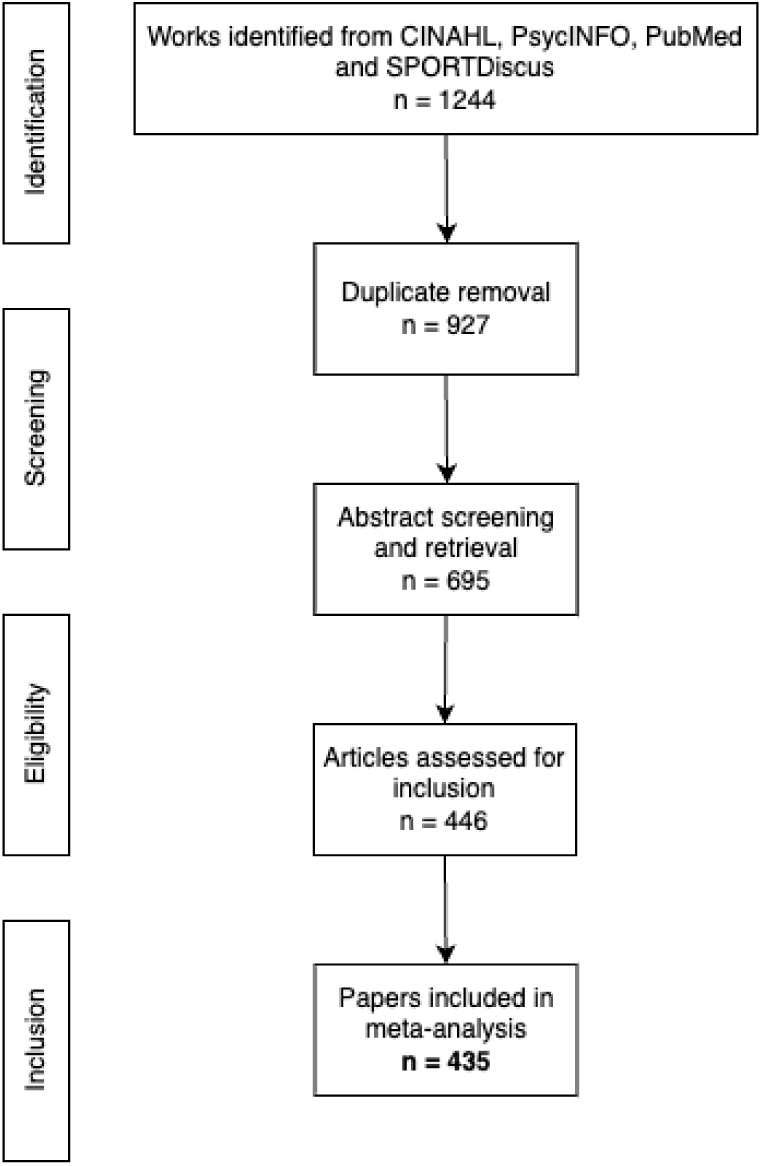
Flowchart detailing paper selection and filtration.

## Results

### Study Characteristics

The total number of participants in the included manuscripts is 3,419,401; due to overlap in datasets between some studies, the number of unique participants is lower. A total of 105 studies used data from a non-unique dataset, the most frequently used datasets included the National Health and Nutrition Examination Survey (NHANES) in 21 manuscripts, the Health and Retirement Study (HRS) in 19, the Dunedin Cohort in 9, the Normative Aging Study (NAS) population in 6, and the Lothian Birth Cohorts of 1921 and/or 1936 in 7. Forty studies used data from multiple datasets. The remaining studies used data unique to those publications.

The included studies focussed on different parameters of their examined populations.

Some studies selected participants based on their age, with many studies (117) considering middle-aged and older adults. Twelve studies focused on young adults, 8 studies included young and middle-aged adults, 68 studies focused solely on middle-aged adults, and 6 studies included long-lived individuals including centenarians and their spouses and family members. Studies were often focused on health outcomes, with 24 studies including individuals with mental health and cognitive conditions, 54 studies focusing on specific physical health conditions including but not limited to chronic cardiovascular disease, ischemic stroke, or infertility. Sixteen studies included cancer patients and survivors, and 9 studies examined HIV positive individuals. Seventeen studies examined populations specific to lifestyle parameters, including 7 which focused on athletes, 2 on tobacco smokers, and a variety of additional studies which examined other lifestyle features. Career choice influence on biological age was examined in 13 studies, while 13 studies examined differential aging rates in twins. Some studies specifically recruited by gender or sex, including 42 studies involving females only, and 26 studies with only males. A plurality of the included studies examined the general population, rather than focusing on subpopulations chosen for a given parameter.

Publication of works on biological aging increased steadily during the period of time reviewed (figure 2). All manuscripts reviewed included some assessment of biological age, sometimes multiple metrics, with telomere length employed by approximately a third of the studies, 162 in total. CpG methylation-based epigenetic clocks were used 226 times in these studies, although some publications made use of multiple CpG-based clocks. A number of published studies describe clocks developed based on novel data sources, such as locomotor activity, facial photographs, and blood biomarkers. Besides these established metrics for biological age, other measures including mitochondrial DNA copy number, transcriptome levels, genomic methylation levels and allostatic load are represented in the papers reviewed here. Figure 3 describes the breakdown of analyses employed in papers published per year.

**Fig 2.**
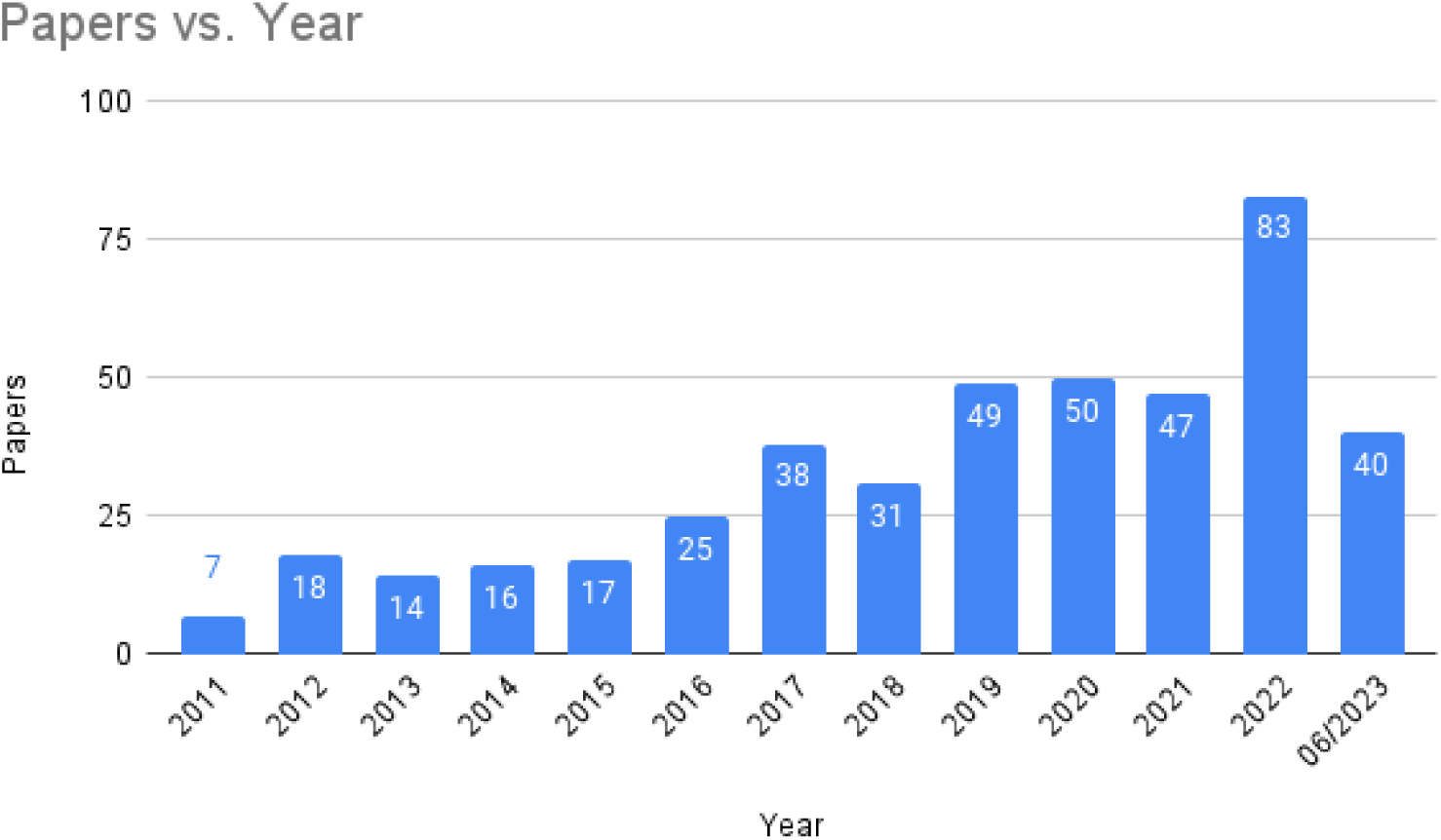
Papers published per year analyzed in this review.

**Fig 3.**
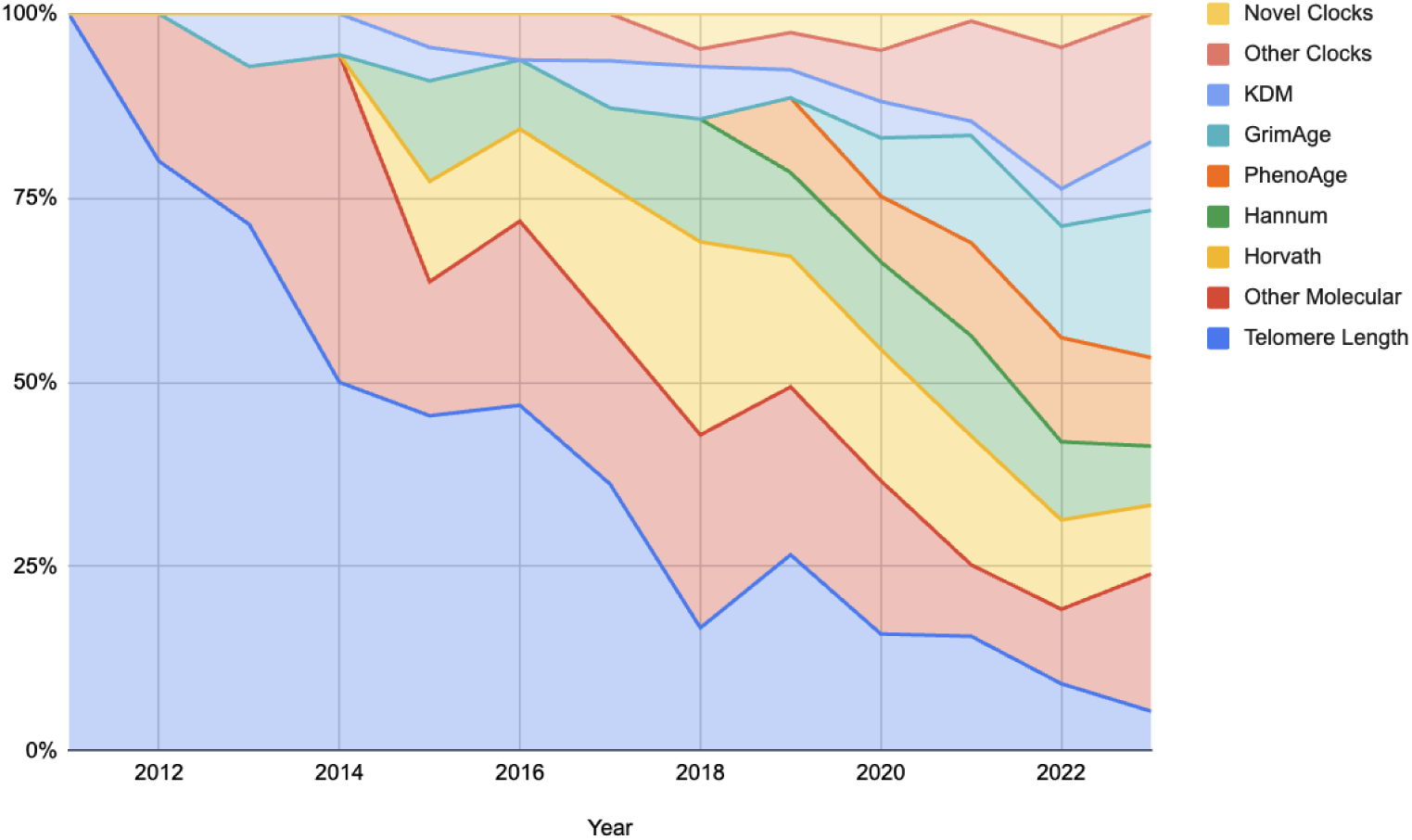
Types of biological aging analyses conducted per year as a percentage of works published in that year.

Overall, studies included were conducted in 43 countries, with the most frequent being the United States (164 manuscripts), followed by 27 in the United Kingdom, and 26 in China. Twenty-three studies occurred in multiple countries, and 4 studies did not report the country they occurred in. Figure 4 details the breakdown by country for those works included in this manuscript.

**Fig 4.**
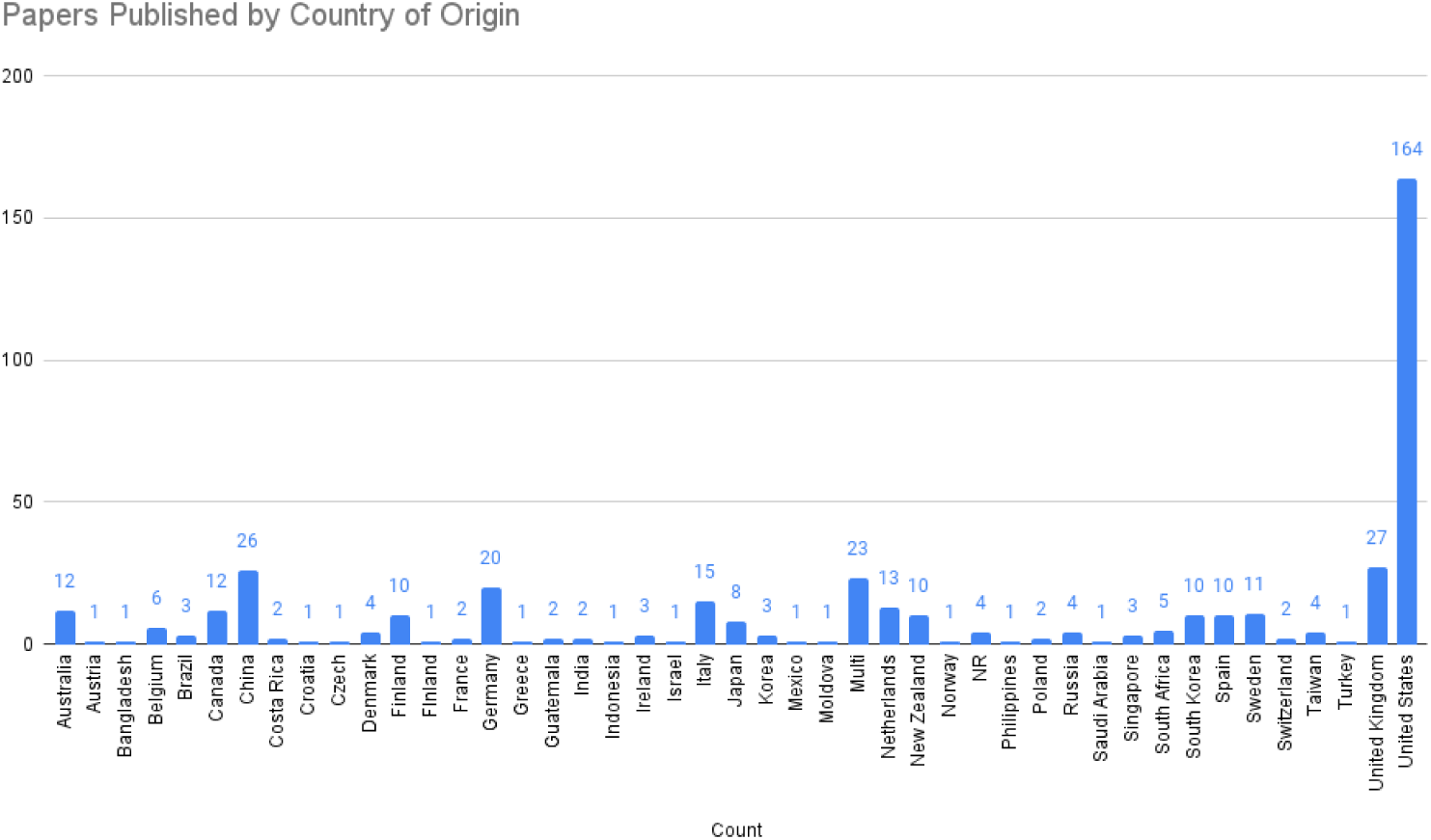
Country of origin for studies included in this review.

### Molecular Biology Features

#### Mitochondrial DNA Copy Number

Mitochondrial DNA (mtDNA) copy number and consequently mitochondrial function are both thought to decrease progressively with age. A study conducted with participants from the NAS observe an association between decreased mtDNA copy number and chronological age, as well as with biological age as determined by the PhenoAge epigenetic clock and with decreased leukocyte telomere length in a follow-up visit [95]. In a 2017 study on the effects of fine particulate air pollution exposure on 552 men enrolled in the NAS, Nwanaji-Enwerem and colleagues found that while mtDNA copy number did not significantly associate with chronological age, it was significantly negatively correlated with Horvath’s pan-tissue clock [293]. Similarly, a pair of studies derived from the VITamin D and OmegA-3 TriaL-Depression Endpoint Prevention (VITAL-DEP) found no association between mtDNA copy number and chronological age [419, 420]. However, a paper published by Praveen *et al.* describe a significant negative correlation between mtDNA copy number and chronological age, with mtDNA copy number higher in women than men among study participants younger than 60 years of age [317]. Furthermore, they found significant positive associations between mtDNA copy number and telomere length, with plasma folate levels, and with vitamin B12 levels among participants older than 60 years of age. A study conducted with participants from the HRS reports a modest association between increased difficulties in activities of daily life and mtDNA copy number [77]. Studies conducted in 2022 report a positive association between mtDNA copy number and healthy lifestyle, a negative association with suicidal ideation and psychological distress, but no significant association with adverse childhood experiences [152, 170].

#### Gene Expression

Gene expression levels are known to vary with age, and some studies have sought to explore the relationship with aging and specific or general gene expression. A study conducted with the African American participants of the Family and Community Health Study used the Peters transcriptome index to determine transcriptomic age acceleration and found an association with histories of both juvenile and adult social adversity and weakly, with male sex, and increased transcriptomic aging [314, 372]. Gheorghe and colleagues analyzed six microarray datasets interrogating three tissues and found distinct time points in each tissue where widespread changes in gene expression occur [133]. Validating previous findings established in the InCHIANTI study population, Holly *et al.* describe a set of five genes, LRRN3, GRAP, CCR6, VAMP5 and CD27, which were similarly identified in the San Antonio Family Heart

Study population, three of which (LRRN3, CCR6 and CD27) were additionally validated in the Exeter 10000 study population [161]. In a pair of sequential studies, first analyzing the Framingham Offspring cohort and following up with the addition of the Framingham Generation 3 cohort and the Rotterdam Study, 448 and 481 genes respectively were identified as associated with a difference between biological age as calculated by the Klemera-Doubal method and chronological age [231, 232]. Other studies have focused on specific genes rather than examining the transcriptome collectively. A 2016 study describes upregulation of the genes telomerase (TERT) and tripeptidyl peptidase I (TPP1) was associated with longer telomeres in endurance athletes [91]. Three related studies investigating ocular health in HIV prevention trial participants and HIV positive individuals found that in HIV negative individuals, expression of CDKN2A was correlated with chronological age, but this was not the case in HIV positive individuals [308–310]. Rentscher and colleagues also examined expression of CDKN2A, but found its expression was not correlated with chronological age, but rather with chronic stress exposure, perceived stress, and accumulated daily stress [331, 332]. In contrast a study of CDKN2A and Nrf2 gene expression found that CDKN2A expression was statistically significantly correlated with chronological age, while Nrf2 was not, in a population of late-stage renal disease patients [391].

Histidine-rich glycoprotein (HRG) expression was found to be positively associated with increased mortality risk and may represent an easily measured indicator of biological aging progression [162]. A study of pathogenic *de novo* DNA methyltransferase 3A (DNMT3A) mutations found affected patients exhibited accelerated aging, with different rates of accelerated aging observed with different mutations of DNMT3A [174].

##### MicroRNAs

MicroRNAs (miRNAs) can epigenetically regulate translation of genes by binding the untranslated regions of their target mRNAs. In a 2014 study of 20 breast cancer patients, Hatse and colleagues identified 15 miRNAs that were differentially expressed between younger and older study participants [151]. In a cell culture study, it was found that rotenone-induced cellular stress could upregulate expression of mir-663, and that cells derived from older donors showed higher levels of induction than those derived from younger donors [422]. The miRNAs mir-21-5p, associated with the inflammatory response, and mir-126-3p, associated with angiogenesis, were both shown to increase with chronological age except in ultracentennarians studied [1].

##### Polygenic Disease Risk

In a 2020 paper, Crimmins and colleagues associate the relationship between polygenic disease risk scores (PGS) and various outcomes, using those risk scores as a proxy for assessing biological aging of specific physiological systems. They find that among non-Hispanic white HRS participants that there is an inverse significant association between the longevity PGS and overall mortality, and an inverse significant association between the general cognition PGS and cognitive dysfunction outcomes [77]. In a study combining data from the Women’s Health Initiative (WHI), Levine and colleagues identified a single nucleotide polymorphism (SNP) that related to both age at menopause and age acceleration by Horvath’s clock [221]. That work found that the minor allele of SNP rs11668344 is associated with earlier age at menopause and increased age acceleration.

#### Non-Clock and Genome-Wide Methylation

##### Non-Clock CpG Sites

In addition to those methylated CpGs that are employed by various epigenetic clocks, CpG methylation globally and at specific loci are also studied in conjunction with biological aging. In 2012, the first paper identifying CpG methylation within the gene ELOVL2 as age-associated was published, finding an increase in methylation in association with increasing chronological age [128]. A study of methylation in the human rRNA locus found one CpG to significantly decrease in methylation with chronological age, while an additional CpG site was found to be associated with impaired cognitive performance and a higher mortality risk after adjusting for age and gender [83]. The authors followed up with a more expansive study in 2019, finding hypermethylation of a CpG contained within the RAB32 gene, and hypomethylation of a CpG contained within the RHOT2 gene, were associated with increasing chronological age [84]. More recently, in a set of post-COVID19 infection and COVID19-free individuals, biological age as determined by a set of four CpGs contained within the genes ASPA, EDARADD, ELOVL2 and PDE4C found that the post-COVID19 infection participants had a biological age 9 years older than their uninfected counterparts, and that this effect was more significant in younger individuals [26, 283]. Kim and colleagues conducted a twin study, looking for CpGs associated with increased frailty in the elderly, and identified a CpG contained within the gene PCDHGA3, as well as several regions containing multiple CpGs, that were associated [189]. A 2015 paper describing the pattern of CpG-reported tobacco and alcohol consumption in conjunction with biological age as determined by Hannum’s clock found that increased cigarette smoking leads to an asymptotic increase in biological aging acceleration, while alcohol consumption showed increased biological aging at the very low and very high extremes of alcohol consumption, with modest age deceleration at moderate consumption levels [23]. In a study of the effects of a relaxation programme on biological age between healthy and myocardial infarction survivor participants, methylation of the genes ELOVL2, C1orf132, KLF14, TRIM59 and FHL2 were measured, and showed that for only the healthy participants that these chronological age-correlated genes’ methylation state reverted to a younger state [311]. Ross and colleagues observed the association of methylation of PAI-1 associated with chronological age, and that biological age measured by that metric decreases over the course of pregnancy; they further found that pregnancy-related increases in body mass index (BMI) led to post-birth decreases in biological age, but that absence of a decrease in BMI in the post-birth year was associated with higher biological age [343].

In a cross-sectional cohort focusing on the most and least socioeconomically deprived individuals in the Psychological, Social, and Biological Determinants of Ill Health (pSoBid) study found that global DNA methylation content was significantly negatively correlated with inorganic phosphate levels, which are in turn associated with poorer diet, lower socioeconomic status and increased cellular inflammatory status [268]. Using Illumina Infinium MethylationEPIC chips in an epigenome-wide association study (EWAS), Tajuddin *et al.* reveal an unexpectedly wide variety of CpGs differentiallymethylated in African Americans relative to white subjects [394]. 4930 CpGs were found to be differentially methylated in African Americans compared to 469 found in European ancestry white participants of the Healthy Aging in Neighborhoods of Diversity across the Life Span (HANDLS) study. Only 301 of those CpGs were identified in common between the two racial groups, and this study identified novel differentially methylated CpGs in both racial groups. A study of methylation at CpG sites in the genes aspartoacylase (ASPA), integrin alpha-IIb (ITGA2B) and cAMP-specific 3’,5’-cyclic phosphodiesterase 4C (PDE4C) found that following 12 months of vitamin B supplementation, ASPA CpG methylation, which typically declines with age, was significantly higher than in control subjects, while PDE4C, which normally increases with age, trended even higher following vitamin supplementation [297]. And in a study on the Lothian Birth Cohort of 1921, Marioni and colleagues identified a CpG highly correlated with facial aging, along with an additional 32 CpGs correlated with facial age and increased mortality risk [257].

##### Genome-Wide DNA Methylation

DNA methylation at sites not associated with established epigenetic clocks and global levels of genomic DNA methylation may be associated with biological aging. A study of genome-wide average methylation in a group of 479 healthy women found no association with chronological age or other methylation-based epigenetic clock measures of biological age, but did find a negative association with the number of live births, and a positive association with age at first live birth among participants [64]. Global DNA hypomethylation was associated with increasing inflammatory status and socioeconomic deprivation, particularly in males [268]. In a more specific study of nine genes, it was determined that seven of those genes showed aging-related patterns of methylation: methylation of the genes GCR, iNOS and TLR2 decreased, while methylation of IFN*γ*, F3, CRAT and OGG increased with increasing chronological age in a study of NAS participants [248].

#### Telomere Length and Attrition

Telomere length as a metric of biological aging is a commonly measured feature, and in the course of this review we collected 162 papers published within a twelve and a half year window that measured telomere length as a proxy for biological age. The majority of the papers (76% or 123 publications) report a significant negative relationship between telomere length and chronological age. Of the remainder, (15% or 25 publications) found no statistically significant relationship between telomere length and chronological age; a minority (9% or 14 publications) did not explicitly characterize the relationship. Telomere length was used to assess the biological aging impact of a variety of variables, including demographic, physical, psychological, clinical, environmental, and social effects.

##### Demographic Parameters and Telomere Length

Demographic variables influence telomere length, including gender and ethnicity. Most publications report longer telomeres observed in female rather than male subjects [3, 18, 54, 56, 71, 132, 164, 185, 234, 277, 408, 416] but a minority of publications reported the inverse, including studies of the Lothian Cohort of 1936 and the Berlin Aging Study II (BASE II) population [15, 275, 280, 410]. A study of 4053 older Chinese adults found that telomeres can lengthen with time, and that this effect was most pronounced in those who exhibited the shortest telomeres at the beginning of the study [169]. Race and ethnicity also were associated with differing telomere lengths. In many studies, African Americans were found to have longer telomeres than did non-Hispanic white subjects [8, 186, 234, 277, 288, 312]. However, von Kanel and colleagues describe shorter telomeres in black South Africans as compared to white South Africans [418]. Few studies reported on the effects of Hispanic ancestry, where individuals with Hispanic ancestry exhibited shorter telomeres than did non-Hispanic whites in an American study [56], but the longevity-exhibiting Nicoyans of Costa Rica exhibited telomeres on average 81 base pairs longer than Costa Ricans from other regions [329]. Family history was also associated with telomere length: men whose mothers were exceptionally long-lived exhibit longer telomeres, while other studies suggest a heritability for telomere length of roughly 55% [164, 186, 187].

##### Physical Parameters and Telomere Length

Numerous studies have been conducted investigating the relationship between an individual’s physical features or biomarkers and biological aging as assessed by telomere length. One frequently studied feature is body weight, typically in the context of BMI, and document the association between shorter telomeres and an elevated BMI [3, 8, 57, 191, 259, 276, 277, 364, 385, 417, 418]. At birth, telomere length was found to be positively associated with birth weight in males, however [204]. Elevated waist circumference was also identified in association with shortened telomeres, as was increased body fat [2, 207, 334, 365]. Inversely, higher lean body mass or appendicular lean mass was associated with longer telomeres [65, 280]. Weight loss reveals an association with elongation of telomeres in female subjects only in a study of bioenteric intragastric balloon recipients, but not in a study considering exercise, metformin, or combination treatment in overweight women [59, 296]. Elevated athleticism is associated with improved telomere retention with aging: studies of master athletes, elite athletes, endurance athletes, cross-country ski racers, ultra-marathon runners and ultra-trail runners confirm this [2, 40, 90, 91, 154, 286, 305, 357]. Conversely, Normando *et al.* report in a Brazilian study that decreased physical activity was associated with longer telomeres in female participants, with the reverse reported for male participants [291]. In many studies, however, regular physical activity was positively associated with telomere length [54, 56, 160, 185, 191]. On the opposite end of the spectrum, the presence of attributes indicative of physical frailty yield inconsistent results [11, 44, 47, 308, 412]. Increased fatigability was also associated with decreased telomere length [185].

Dietary considerations also play into telomere length. A study using data from the Korean Genome Epidemiology Study population found that a traditional Korean diet including seafood, whole grains, legumes, vegetables and seaweed was associated with longer telomeres as compared to a Western diet emphasizing red and processed meat and refined grains [208]. Among hypertensive patients, very low and moderate intake of dietary copper was associated with decreased telomere length, a finding thought to be due to the antioxidant properties of copper [137]. Elevated serum levels of inorganic phosphate, associated with poorer diet, or a poor diet score, were similarly associated with shorter telomeres [268, 363, 446]. Higher levels of vitamin C and potassium were associated with longer telomeres in a study of older Koreans [209]. Analyses of NHANES study data find that consumption of lower fat percentage milk, higher dietary fiber, greater quantities of fruits and vegetables consumed and lower blood levels of gamma-tocopherol (although not supplementation with vitamin E) are all associated with longer telomeres [402–405]. Alcohol consumption and alcohol abuse are associated with shorter telomeres [210, 364, 386, 417, 418], as is cigarette use [54, 56, 104, 186, 363, 385, 399, 429] and illicit drug abuse [51, 233].

In terms of genetics, the association of at least two SNPs have been associated with telomere length: possessing the alternative allele for rs4880, in the superoxide dismutase 2 (SOD2) gene, was associated with shorter telomeres, as was possession of the alternate allele for rs9939609 or rs2736100 [154, 336, 365]. Carriers of a mutant allele of the SERPINE1 gene exhibited longer than wildtype telomeres [187].

Blood-based biomarkers are also associated with telomere length. Elevated HDL cholesterol is associated with longer telomeres [249, 334] although at least one study reports a reverse relationship [398]. Elevated triglycerides and glucose both associate with decreased telomere length [398], as do elevated C-reactive protein, IL-6 and TNF*α*, particularly when the latter two are both elevated [299, 333]. In studies of Australian men, Yeap and colleagues observed that circulating estradiol, IGF1, IGFBP3 and dihydrotestosterone levels are all associated with increased telomere length, while sex hormone binding globulin (SHBG) levels are negatively correlated [449–451]. A later study expanded upon this finding, noting that telomere length and low testosterone association was mediated by SHBG levels [258]. Particularly in older adults, high plasma ferritin levels are associated with shorter telomeres, while elevated folate and vitamin B12 levels correlate with longer telomeres [237, 317]. In two published studies, Maeda and colleagues observe gender-specific associations with specific blood composition features and telomere length: red blood cell counts were positively correlated with lengthened telomeres in Japanese study participants of both genders, while other blood parameters differed [249, 251]. In women, elevated urinary malondialdehyde levels, an indicator of oxidative stress, was positively associated with telomere length while urinary cortisol was negatively associated [20, 104]. Finally, gut flora composition was found to be associated with methylated telomere length but not unmethylated telomere length [252].

##### Psychological Parameters and Telomere Length

Mental health and psychological features are known to have an impact on telomere length. In a study of 468 American military veterans, Watkins and colleagues found that greater hostility and difficulties controlling anger were significantly associated with decreased telomere length [429]. Research analyzing the experiences of 600 adults with mild to moderate cognitive impairment found that those who experienced physical, psychological, or especially combination abuse exhibited significantly shorter telomeres [108]. Among a study of healthy middle-aged women, shorter telomeres were associated with an increased propensity for mind wandering, but not perceived stress or rumination [103]. Work-related stress has been investigated, finding longer telomeres among those who report less work-related exhaustion and greater learning opportunity at work, and shorter among those who report the effects of depersonalization (“burnout”), decreased social support, and increased reliance on compensation behaviours in the workplace [3, 68, 432]. However, other studies did not confirm the relationship between chronic stress exposure, perceived stress, and accumulated daily stress with telomere length [331]. Results regarding the association between telomere length and major depressive disorder are mixed, with some studies finding a negative association [276, 408, 438] or a negative association only in conjunction with other history such as childhood physical neglect or racial discrimination [61, 210, 234, 414], or no significant association [152]. No association between telomere length and autism diagnosis or hypersexual disorder diagnosis were detected in a pair of studies published in 2022 [43, 301]. In the Strong Heart Family Study, focused on the health of Indigenous Americans, men with severe depressive symptoms were found to have shorter telomeres, but not men with less severe symptoms or women with any severity of depressive symptoms, while the Dunedin Study found men, but not women, with internalizing disorder and depression exhibited shorter telomeres [360, 465]. Other studies have explicitly identified no relationship between the presence of depression and an impact on telomere length [345]. The presence of anxiety was associated with impaired telomere length [160, 333, 360].

Childhood adverse events, including physical, psychological or sexual abuse, may induce a change in telomere length, but some studies demonstrated the absence of an association [170]. In a study of Finns who were separated from their parents during the Second World War as children, it was found that those individuals separated from both parents who also experienced physical or psychological abuse exhibited shorter telomeres than those participants without those experiences, decades later [350]. Mason *et al*. found no association between telomere length and a history of childhood abuse in a study of 1135 women recruited for the Nurses’ Health Study II [263]. Contrary to expectations, former Swiss indentured child servants who exhibited PTSD had longer telomeres than control individuals [202]. In other studies, PTSD was associated with shorter telomeres, and increasingly shorter telomeres for those individuals whose PTSD was coincident with an experience of childhood abuse [300].

Mental health interventions may offer some remedy: a study of the effects of meditation practices found that lovingkindness, but not mindfulness meditation, was associated with reduced telomere loss over the course of the trial [218]. In a study of myocardial infarction (MI) patients and control individuals, sixty days of relaxation practice preserved telomere length in controls only [311]. Lastly, sleep and sleep quality has been assessed in conjunction with telomere length. Work done with 672 participants of the Multiethnic Study of Atherosclerosis (MESA) found associations between reduced telomere length and severe obstructive sleep apnea and other sleep parameters [56]. A study of 887 Korean participants in the Korean Genome Epidemiology study found that regardless of the presence of apnea, time spent snoring during sleep was associated with decreased telomere length [364]. Certain prescription sleep aids were associated with a shortening of telomeres among men [253]. Delayed circadian rhythm, self-reported moderately late chronotype, chronic short sleep and later sleep onset time were all associated with shorter telomeres, while self-reported extremely early chronotype was associated with longer telomeres, in a study of 2936 Dutch adults [441].

##### Clinical Parameters and Telomere Length

States of health and disease have shown various associations with telomere length. In terms of reproductive health, telomere length was significantly positively associated with the number of surviving offspring a woman has, and in a further study, experience of child mortality was significantly negatively associated [19, 20]. Smeets and colleagues found that there is a positive correlation between gestational age and telomere length in young adults [376]. Telomere length was associated with maternal energy reserves as represented by arm fat area [204]. No link with pre-eclampsia was detected for telomere length in a 2022 study, nor was a link detected with early versus normal ovarian aging in a separate study [69, 319].

Overall risk of mortality typically correlates with decreased telomere length, although not reported by all studies [125, 135, 163, 228]. Diabetes and diabetes duration are associated with telomere loss [4, 54, 67, 210, 398, 406]. Conversely, diabetes interventions were found to lengthen telomeres in participants, particularly among those who began the intervention program with the shortest telomeres [168]. Similarly cardiovascular disease and cardiovascular parameters, including retinal microvasculature, are generally associated with telomere length [67, 90, 91, 144, 149, 165, 185, 250, 276, 305, 327, 333, 369, 460]. In a study of patients implanted with cardioverter defibrillators (ICDs), those who had received therapy from their ICDs had reduced telomere length than did those who had not received ICD therapeutic intervention, and that this feature was highly predictive of ventricular arrhythmia events [351]. A study of Italian early onset acute myocardial infarction (MI) patients revealed an association with hypertensive status and reduced telomere length but did not associate early onset MI with reduced telomere length [344]. Among chronic obstructive pulmonary disease (COPD) patients, telomere length was not associated with spirometric lung function measurements, but was found to be in association with the six minute walk distance and mean daily step count measures [425]. A small study of spinal surgery patients found that those patients with the shortest telomeres were most at risk for post-operative complications than those patients with longer telomeres [346].

Diagnosis or a history of cancer was associated with shortened telomeres, and colorectal tumour tissues exhibit shorter telomeres than adjacent cancer-free tissue [8, 125, 185, 267]. A study of early breast cancer patients found that telomeres were not shorted by twelve months’ chemotherapeutic treatment [48]. Infection with HIV was associated with a reduction in telomere length [276, 281, 309, 310, 417, 418]. In a study of 88 women, 66 of whom were diagnosed with fibromyalgia, it was found that telomeres were shortest among those fibromyalgia patients who exhibited higher pain and more severe depressive symptoms [148]. Multiple sclerosis patients were found to have shorter telomeres than age-matched controls but did not exhibit an accelerated loss relative to controls [142]. A study of schizophrenia patients found that while there was no significant impact to telomere length based on diagnosis alone, there was an interaction between gender and diagnosis in which schizophrenic women showed the greatest reduction in telomere length relative to healthy controls and schizophrenic men [440].

Dhillon and colleagues reveal reduced telomere length among carriers of the APOE-*ε*4 allele compared to non-carriers, and an inverse relationship between plasma levels of advanced glycosylation end products and telomere length [92]. Other findings related to telomere length and Alzheimer’s disease and dementia are mixed: a 2020 study found that Alzheimer’s disease patients exhibited accelerated telomere loss relative to individuals with mild cognitive impairment (MCI), contradicting a previous 2012 study that found that MCI patients exhibited shorter telomeres than controls and Alzheimer’s disease patients, and that those MCI patients who later progressed to Alzheimer’s disease exhibited longer telomeres than those who did not progress [211, 284]. Work with the Washington Heights-Inwood Community Aging Project population found that in women, telomeres were shorter in those with dementia and were also a risk factor for developing dementia [163]. Related, a 2022 study describes an association between elevated aging in the brain and shorter telomere length [455]. A study of elderly adults without dementia found shorter telomeres were associated with greater degree of subcortical atrophy as detected by magnetic resonance imaging (MRI) [436]. While not associated with rheumatoid arthritis (RA) severity, a study including 145 RA patients and 87 controls found that reduced telomere length was associated with a worse coronary artery calcium score in RA patients but not controls [302]. Patients with Prader-Willi syndrome (PWS) exhibit lower median telomere length than both controls and non-PWS participants born short for gestational age and subsequently treated with growth hormone [96].

##### Environmental Parameters and Telomere Length

Features such as living environment as well as exposure to various toxic chemicals may influence telomere length. Satisfaction with one’s neighbourhood correlates positively with telomere length [132]. In a study examining the KORA F4 and NAS cohorts, it was found that air pollution components had gender-related effects, where only men showed telomere shortening in response to increasing black carbon particulate exposure [428]. An additional study on air pollution’s effects on telomere length found that for males, mid-adulthood exposure was especially impactful, while for females, air pollution exposure levels perinatally were more critical [18]. Exposure to an organophosphate insecticide caused shortened telomeres among those in the second quartile of exposure, in a study of 1724 NHANES participants [298]. Data derived from the Scottish Family Health Study found that increasing exposure to second hand cigarette smoke caused increasing telomere attrition [244]. An inverse association between telomere length and total dietary exposure to persistent organic pollutants and polychlorinated biphenyls (PCBs) was identified in a 2022 study [5]. Among women, higher phthalate and phenol metabolites in urine was associated with both an increased risk of breast cancer and with telomere attrition, with telomere length being a modifier in the association between exposure and breast cancer risk [463].

##### Social Parameters and Telomere Length

Socioeconomic status is frequently studied in conjunction with telomere length. Higher household income is positively associated with telomere length [61, 363, 452] as is higher educational attainment [132, 164, 191, 392], marriage [452] and home ownership (as opposed to renting) [54]. In a study of Detroit residents, it was found that the effects of poverty led to greater telomere attrition in white participants than in African American participants [132]. However, social class was not associated with telomere length [392] or unexpectedly, was found to be associated negatively with telomere length in men [453]. The West of Scotland Twenty-07 Study population was examined in a study finding that members of different generations exhibited different patterns of response to various socioeconomic parameters [338]. Parental effects are observed as well, where a parent’s occupation or educational attainment influenced the telomere length of their offspring [54, 177, 291]. Reports of lifetime discrimination, including years after the incidents in question, were significantly associated with decreases in telomere length [58]. Other influences on telomere length included a high belief in justice for the self being associated with reduced loss of telomeres [247] and unfair treatment to self being associated with increased loss of telomeres [330]. Finally, telomere length was associated with land use mixture increase, particularly among foreign-born and less acculturated Mexican Americans [466].

#### Other Molecular Biomarkers

##### Non-Genetic Biomarkers

Use of non-genetic markers, such as serum protein or metabolite levels, and the use of physiological features such as forced expiratory volume or systolic blood pressure, are often used in conjunction to assess biological age.

Studying middle-aged adults and beginning with a collection of 21 blood biomarkers, Levine found that a set of merely seven: C-reactive protein (CRP), glycated hemoglobin (HbA1C), serum alkaline phosphatase, forced expiratory volume (FEV1), systolic blood pressure and in men, cytomegalovirus optical density and serum albumin while in women, total cholesterol and serum urea nitrogen, were highly informative with regards to biological age, and produced significantly better mortality prediction than chronological age or other biomarker combinations tested [219]. Beam and colleagues use these markers in their 2020 work, where they demonstrate that in a study of middle-aged twins, biological age significantly negatively correlates with physical functioning and certain aspects of memory [25]. In the same year Crimmins describes work on the HRS population, finding the same markers associated with increased difficulty in conducting activities of daily life, multimorbidity, cognitive dysfunction and mortality [77]. These markers were also used in a study that found that African Americans exhibited an elevated biological age by this measure, and that the risk of mortality was greater in African Americans relative to white study participants, revealing an underlying difference in the rate of biological aging between racial groups [220]. Using a subset of these markers, it was found that the rate of biological aging is changing over time: while biological age is lower for younger subpopulations, improvements in biological age deceleration are greatest among the oldest male study participants, possibly due to cultural shifts in health modifications including smoking cessation [222]. Studying a further restricted subset of these markers in a large merged data set found that 481 genes significantly change their expression levels in a manner dependent on this biological age measure, with most of the genes exhibiting a positive association with biological age [232]. The authors speculate that this may indicate that changes in biomarkers may be a consequence rather than a contributor to biological aging. In a study employing Levine’s ten markers plus twelve additional biomarkers, a 2023 study identified accelerated biological aging in non-Hispanic Blacks and Hispanics relative to non-Hispanic whites regardless of sex, that was modified by educational attainment [109].

An overlapping group of clinical biomarkers was identified in research derived from the Canadian Study on Health and Aging population, where serum albumin, calcium, serum creatinine, diastolic blood pressure, hemoglobin, serum alkaline phosphatase, inorganic phosphorus, total protein, thyroid-stimulating hormone (TSH) and blood urea nitrogen were found to be significantly correlated with chronological age [282]. The same set of biomarkers were used in a study combining the Rotterdam Study and NHANES study populations and found that biological age as measured by these markers was a better predictor of diabetes, stroke, cancer and mortality than chronological age, and roughly equivalent to chronological age at assessing the risk of coronary heart disease and all-cause mortality [430]. In an investigation of the BASE and BASE II study populations, a novel set of biomarkers was found to be more accurate than the Vidal-Bralo or PhenoAge clocks in determining mortality and subjective health [97]. A 2021 paper describes a combination of Levine’s markers along with the Targeted Aging with Metformin (TAME) assay panel and found that these together were good predictors of multimorbidity, but not better than chronological age at predicting cognitive dysfunction [21, 78]. A 2021 study in the HRS population found biological age as determined by Levine’s biomarkers was associated with persistent feelings of loneliness, as well as feelings of social isolation [79]. These same biomarkers were associated with elevated physical activity among COPD patients and improved biological age, as well as with a diet relatively high in carbohydrates with moderate lipid and protein components, associating the marker set with general health [358, 425].

In the Singapore Longitudinal Aging Study population, a study was done that found that 8 and 10 biomarkers adequately described biological aging in men and women respectively, and found that a wide range of lifestyle, behavioural, and socioeconomic factors were linked to the rate of biological aging, including housing type, marital status, physical activity and diet [290]. Using only physiological and physical fitness variables, Jee and colleagues describe a set of 8 partially overlapping parameters for each sex that model biological age in the Korean population, and they describe an association of poor biological age by this measure with both elevated BMI and sarcopenia [172]. Later work with the Korean KNHANES III study examined a set of 31 biomarkers and focused on seven: systolic blood pressure, waist circumference, glutamic oxaloacetic transaminase, ferritin, blood urea nitrogen, creatinine, and FEV1, finding an association of biological age by this metric with both diabetes and increased glucose tolerance [173]. Meisel and colleagues developed a blood-based biomarker panel with significantly better predictive power of tooth loss than chronological age [278]. Examining a panel of 36 blood biomarkers including indicators of glucose metabolism, cardiac and renal function, and common hemochrome markers, a study of 4592 participants found that those with a higher polyphenol antioxidant content diet exhibited a lower difference between chronological and measured biological age, particularly among men [105]. Work on the Northern Sweden Population Health Study cohort examining 144 plasma protein profiles and anthropometric measurements found dietary and lifestyle effects that were associated with changes in biological age, finding consumption of fatty fish and 3 to 6 daily cups of coffee beneficial, while smoking, high soda consumption, obesity and minimal exercise detrimental [102].

In a study of 1581 older adults, a battery of parameters including body composition, parameters of energy availability and consumption, homeostatic equilibrium measures, and neurodegenerative and neuroplastic measures were assessed in relationship with chronological age, finding a substantial number of features that contributed to biological age [203]. A study of biomarkers found a set of 12 metabolites, four of which were linked to the fatty acid/tri-citric acid cycle metabolism, with the remainder related to glycolysis, predicted a biological age significantly correlated with chronological age in a set of 604 older adults [175]. In a study of 47 plasma metabolites, 37 were found to be significantly associated with age, including those involved in bioenergetic pathways, suggesting a decrease in mitochondrial beta-oxidation and accumulation of unsaturated fatty acids leads to an increased in both oxidative damage and inflammation with age [377]. Thirty-four inflammation, vitamin B and kynurenine pathway blood biomarkers were assessed for their biological age predictive capacity; 29 of these were associated with chronological age, 17 were found to be associated with mortality at followup, and a score composed of 10 biomarkers was associated with higher all-cause mortality in the study population [99]. Among 3558 participants in a wellness program, Earls and colleagues found that among clinical laboratory results, metabolite panels, and proteomics data, biological age as derived from proteomics data outperformed the other two data types in terms of correlation with chronological age and mean absolute error [101]. They found a variety of poor health conditions including obesity, hypertension, elevated cholesterol, lung infection, type II diabetes and breast cancer associated with this measure of biological aging, and that participants in the wellness program exhibited a decline in biological age after completion of the program. In a study examining the Moli-Sani study population, an increase in energy-adjusted dietary inflammation index and dietary inflammation score was associated with an increased biological age as estimated by blood biomarker levels [261]. Other efforts to leverage proteomics data for biological age determination include mass spectrometry-based analyses: a measure based on five peptide fragments including apolipoprotein A1 (ApoA1), fibrinogen alpha chain, complement C3, complement C4a, and breast cancer type 2 susceptibility protein in a 2020 publication, while a 2019 study describes mass spectrometry data biomarkers developed in the MARK-AGE study, finding that in HIV patients, biological age was elevated relative to healthy control donors [50, 52, 88]. Other physical parameters assessed include urinary markers. A 2022 study demonstrated both positive and negative correlations with chronological age for urinary oxidative stress markers, suggesting this as a useful metric for calculating biological age [285]. Saliva has also been explored as a source of biological aging data: in a study of 79 saliva proteins, 69 were found to decrease in healthy elderly participants, while 37 proteins increased in Alzheimer’s participants compared to controls [72]. Non-invasive cardiovascular measurements have also been assessed for their informational content: a 2023 study determined that the left atrium reservoir strain, left atrium conduit strain rate and the ratio of left atrium conduit strain rate to booster strain rate was found to be reduced in older adults as compared to younger adults [194].

More specific studies focusing on domains of related proteins have revealed patterns of pathogenesis. Work restricted to measurements of senescence-associated secretory phenotype (SASP) markers found an association between increased frailty and certain SASP markers among a merged cohort of elderly healthy adults, aortic stenosis surgery patients and ovarian cancer patients, with levels of GDF15, OPN and TNFR1 significant among all three subgroups [352]. A previous study had identified GDF15 in particular as positively associated with increased morbidity, cognitive impairment and decreased functional independence, as well as being a significant mortality predictor in a study of older adults [349]. In a study of older adults with late onset depression, it was found that depressed adults exhibited a higher GDF-15 serum level than did non-depressed age matched controls, and that those depressed adults with later onset of depression had elevated GDF-15 as compared to earlier onset [264]. Further, a 2022 study later identified a moderate correlation between the SASP marker GDF15 and chronological age in postmenopausal women, suggesting that SASP markers may be indicative of biological age in that population [366]. An investigation of a panel of 9 inflammatory biomarkers including C-reactive protein and IL-6 was used to develop an inflammatory biological age measure, finding seven of the nine were significantly correlated with chronological age [231]. Furthermore, they identified 448 genes whose expression was significantly associated with inflammatory biological age. A study focused on a restricted set of five biomarkers found an association between CRP and BMI as well as waist circumference; the same study identified a correlation between GDF15 and smoking pack-years in men, but not women [382].

Other studies approached the question of disease-specific, rather than pathway-specific, genes and proteins underlying biological aging. In studies of breast cancer, Brouwers and colleagues found association of IL-6, IGF1, MCP1 and TNF*α* with chronological age, and frailty was particularly associated with levels of IL-6 [47, 48]. Findings of significance for IL-6 and TNF*α* were confirmed in a 2022 paper, with the additional finding linking circulating levels of those proteins to increased multi-morbidity risk [388]. Circulating IL-37 levels were found to be lower in older adults compared to middle aged adults, and an association with the IL-37:CRP ratio with maximal oxygen consumption (*V O*_2_ max) and body mass was found; the IL-37:IL-6 ratio was also positively correlated with VO2 max and lean body mass [49]. Berben and colleagues found combinations of blood biomarkers associated with frailty and tumour immune infiltrate characteristics in a study of older breast cancer patients [32]. Additionally, sampling methods beyond blood draws have been explored: a 2016 study explored urine metabolome data in a study of bariatric surgery patients and controls, finding a metabolic age score that was predictive of survival at 13 years of follow-up, as well as of weight loss for the bariatric surgery patients [155]. Fraszczyk and colleagues, also in a study of bariatric surgery recipients, found an improvement in levels of liver enzymes following surgery, suggesting these markers may also reflect a metabolic biologic age metric [120]. In individuals with mild traumatic brain injury, it was determined that chronological age at injury is associated with increased brain aging following that injury as determined by magnetic resonance imaging (MRI) [7].

Studies of hormone levels have revealed patterns underlying biological aging in different disease states. In a study of COPD patients, dehydroepiandrosterone (DHEA) and growth hormone levels were measured, finding an inverse relationship with chronological age and significant depression of DHEA and growth hormone levels in those patients, suggesting the biological age of a COPD patient is as much as 24 years older than a control subject of the same chronological age [183]. An Indonesian study of fertility-challenged women found a negative correlation between chronological age and antral follicle count and anti-Mullerian hormone, and a positive correlation with follicle stimulating hormone (FSH) and chronological age [439]. Data from the Baltimore Memory Study was employed to identify a relationship between increasing waking cortisol levels and chronological age, as well as lower average waking cortisol levels and slower diurnal decline in African Americans while socioeconomic vulnerability was associated with a faster diurnal decline after adjustment for race/ethnicity [348]. An additional study found morning and evening cortisol secretion was highest in men with severely accelerated (*i.e.*, eight or more years acceleration), suggesting a disruption of circadian rhythm may be a feature of accelerated biological aging [138].

Individual plasma protein levels have also found association with biological aging. A study of N-terminal fragment B-type natriuretic peptide precursor (NT-proBNP) found a significant association with NT-proBNP levels and risk of mortality in multivariate Cox regression analysis [287]. Plasma soluble urokinase plasminogen activator receptor (suPAR) was found to have a positive correlation with age, and elevated suPAR was associated with lower functional capacity, poorer cognitive function and cognitive decline, accelerated facial aging and increased physical limitations [328]. That study also observed a relationship between higher tobacco use or lower physical activity and higher levels of suPAR. suPAR was also identified as elevated in older adults and associated with increased risk of cardiovascular disease [194]. A significant positive association between levels of HbA1C and chronological age was identified in healthy study participants, suggesting that diabetics with elevated HbA1C may exhibit accelerated biological aging [393]. A study of 701 middle-aged Han Chinese found 19 immunoglobulin G (IgG) glycans that were correlated with chronological age, 10 of which were consistently correlated in both men and women [454]. And in a small study of men, terminal galactosylation of ferritin was posited as a biomarker of healthy aging [100].

Macromolecular cellular features, such as the presence of damage foci and damaged telomeres, were found to be positively associated with donor chronological age in a study of cultured fibroblasts [423]. The length and coverage area of elastic fibre within the dermis was found to decrease with increasing chronological age, as well as observed flattening of the dermis, suggesting that dermal characteristics can serve as a measure of biological aging [421]. Related, hair whitening, confirmed to correlate positively with increasing chronological age, was also significantly linked to the presence of coronary artery disease, which allows for the potential of premature hair whitening to be an indicator of advancing biological aging [193].

##### DNA Damage

Accumulated DNA damage represents one aspect of biological aging. In one study of subjective cognitive function in women diagnosed with breast cancer, the authors found a positive association between DNA damage in leukocytes as assessed by the comet assay and chronological age and reduced executive function scores with increasing DNA damage accumulation [57].

##### Allostatic Load

Allostatic load is a measure of accumulated chronic stress and life events on different physiological systems, including cardiovascular function, renal function, immune system status and neuroendocrine systems and represents a degree of ‘wear and tear’ associated biological aging [141]. A study of the Lothian Birth Cohort of 1936 assessing the relationship between brain predicted age differences (brain-PAD) and various measures of biological aging found a positive association between worsening brain-PAD scores and increased allostatic load [71]. Assessing gender-specific relationships with allostatic load, McCrory and colleagues found in a study of 490 older Irish adults that metabolic allostatic load was positively associated with epigenetic age acceleration in men, while cardiovascular and metabolic allostatic load were positively associated with epigenetic aging acceleration in women [269]. A later study found that higher allostatic load associated positively with higher all-cause mortality risk, with worse performance on physical functioning tests, and with lower educational attainment and material resources [150]. In a study of reproductive history among female participants in the NHANES population, premenopausal women showed no association with allostatic load, number of births or time since last birth, while postmenopausal women revealed a positive significant association between the number of births and allostatic load [367]. Finally, allostatic load was found to correlate significantly with other measures of biological aging, including Horvath’s pan-tissue clock, Hannum’s clock, Levine’s PhenoAge clock, homeostatic dysregulation measures, and biological aging as measured by the Klemera-Doubal method [269, 367].

##### Basal Oxidation

Measures of oxidation as an assessment of metabolic rate may be associated with biological aging. A study using plasma levels of the anti-aging protein S-klotho as a marker of youthfulness found that elevated levels of S-klotho were associated with increased basal fat oxidation and decreased basal carbohydrate oxidation, but not associated with chronological age [6]. A 2023 study reports a negative association between klotho levels and dietary inflammatory index [446].

##### Physiological Dysregulation

Physiological dysregulation incorporates data from multiple organ systems to assess an overall level of dysfunction associated with increasing age. In a study of 93 male pilots, Bauer and colleagues measure 18 markers of organ function and find a non-linear relationship between increasing physiological dysregulation until approximately ages 45 through 50, and then a decline in overall physiological dysregulation [22]. A 2022 study characterized patterns of nutrition associated with minimal biological aging as measured by physiological dysregulation, including moderate uptake of macronutrients and ingestion of vitamins E and C, especially in combination [358]. Using 12 biomarkers, a physiological dysregulation-based calculation was able to predict mortality risk in a Chinese population [238]. Among hospitalized COVID-19 patients, a Russian study observed that with increases in physiological dysregulation, the risk of both deterioration and mortality also increased [387].

##### Telomerase Activity

Associated with telomere length is activity of the multi-subunit enzyme telomerase, which acts to restore telomere length lost during DNA synthesis in cell division. Chronological age is generally reported as having an inverse relationship with telomerase activity [57, 111]. In a study of patients who had received implanted cardioverter defibrillators (ICDs) following diagnosis of coronary artery disease and previous myocardial infarction, those patients who had received therapy from their ICDs exhibited elevated telomerase activity relative to those who had not [351]. A study of 94 women who had been diagnosed with breast cancer found that the cognitive parameters executive function, attention, and motor speed were all positively correlated with telomerase activity [57]. Telomerase expression was found to be higher in endurance athletes with the highest cardiorespiratory fitness and the lowest resting heart rates compared to those less fit athletes and controls [91].

##### Molecular Markers of Bone

For forensic or archaeological purposes, biological age of human remains may need to be determined. As blood-based and DNA-based biomarkers may be absent or degraded, methods to correlate biological age with skeletal features have been developed. In a 2017 study of Mexican and Mesoamerican remains, auricular surface (AS) and pubic symphysis (PS) were used to predict biological age of the decedents. Among those remains for whom chronological age was known, AS biological age prediction exhibited a mean difference of 4.1 years (standard deviation 3.0 years) from chronological age, while PS-based prediction achieved a mean difference of 4.5 years (4.2 years) [74].

### Biological Aging Algorithms and Clocks

#### Horvath’s Pan-Tissue Clock

Perhaps the best known among epigenetic clocks is the Horvath pan-tissue clock, developed in 2013 using 51 healthy cell types and tissues and elastic net regression to predict biological age [166]. The Horvath pan-tissue clock calculates biological age based on the methylation status of 353 CpGs and is known for its high correlation with chronological age and its substantial performance across different biopsied tissues and cells. The Horvath pan-tissue clock is a first generation epigenetic clock, based exclusively on DNA methylation and represents a major step forward in the measurement of biological aging. In the 108 papers published in the review window using the Horvath pan-tissue clock that we assessed for this review, among those that reported correlation with chronological age all reported a substantial positive correlation. The Horvath pan-tissue clock also exhibits moderate to high correlation with other clocks, including the Hannum epigenetic clock, the Horvath skin and blood clock, the Weidner clock, and with the second generation clocks Dunedin Pace of Aging, PhenoAge and GrimAge [29, 31, 64, 82, 98, 107, 228, 269, 294, 296, 340, 342, 379]. This clock also correlates negatively with mtDNA copy number [95]. However, the Horvath pan-tissue clock does not appear to be correlated with measures of facial aging [29, 257]. There may be a gender specific component, as some reports include findings of a faster Horvath clock-based epigenetic aging observed in men than in women [17, 18, 36, 37, 98, 110, 181, 204, 235, 269, 303, 390]. One report detailed the reverse finding [260]. Ethnicity may also play a role: epigenetic aging by Horvath’s clock was found to be accelerated in white versus African American study participants, and versus Hispanic study participants, but these findings were not universally reported [37, 110, 139, 394]. The rate of aging by this clock may be partially heritable, as a study of twins and non-twin sister pairs found that monozygotic twins had greater correlation of deltaAge, the measure of chronological age less biological age, than did dizygotic twins or non-twin sibling pairs, and a study of 4658 elderly adults estimated the heritability of Horvath deltaAge at 43% [225, 256].

As the Horvath pan-tissue clock was partially trained on cultured cells, it has also been tested on cell lines, and in an experiment involving long-term cultured fibroblasts found that epigenetic age as measured by the pan-tissue clock accumulated as the cells grew but ceased to increase once cells reached senescence [389]. While the Horvath pan-tissue clock was developed while the Illumina Infinium Human Methylation 450k BeadChip was predominant, a newer chip, the Infinium MethylationEPIC BeadChip later became available, boasting an increased 850,000 CpG sites interrogated. A 2018 study found that, despite the absence of 19 of the 353 CpGs of the Horvath pan-tissue clock, MethylationEPIC BeadChips did not suffer from decreased performance in epigenetic age prediction [272]. Studies of the accuracy of the Horvath pan-tissue clock has found a tendency to underestimate middle-aged and older individuals: a study of middle-aged twins found a systematic underestimation of age relative to chronological age, and cortex samples biopsied from individuals greater than 60 years of age were similarly underestimated [368, 383].

Biological age as measured by the Horvath pan-tissue clock has been repeatedly associated with mortality risk. In a study of 378 twins, it was found that among the oldest twins studied, the twin with the oldest biological age by Horvath’s clock was the first twin to die in 69% of cases, which corresponded to a more than three times the risk of mortality as compared to their twin [70]. In a study including the Lothian Cohorts of 1921 and 1936, a five year increase in deltaAge corresponded to an increase in mortality risk of 9% in a model adjusted for multiple factors [256]. A 2020 study found that a one standard deviation increase in biological age resulted in a 17% increase to mortality risk, a finding that was further elevated in ever smokers [228]. This correspondence with mortality risk was not always observed: Gao and colleagues developed a mortality risk score that did not associate with Horvath clock age acceleration, while a study involving participants from the Louisiana Healthy Aging Study cohort found that frailty index was a better predictor of mortality than both chronological age and Horvath biological age, and a further study found no sensitivity to age-related decline in clinical parameters [125, 188, 270]. An additional study considering the Lothian Birth Cohort of 1936 found that Horvath’s biological age in conjunction with brain predicted age difference, based on neuroimaging assessments, was a better predictor of mortality than either individual measure [71]. Morbidity is also predicted by Horvath’s biological age: elevated epigenetic age was associated with multimorbidity and was associated with greater chronic, but other studies found no association with physical frailty [77, 80, 122, 123]. However, a 2022 study of postmenopausal women found participants less likely to achieve age 90 with intact mobility as their Horvath’s age acceleration increased [171]. In a study of young adults and nonagenarians, cytomegalovirus (CMV) seropositivity was associated with elevated Horvath age in both groups [178].

Beyond mortality and morbidity, Horvath pan-tissue age acceleration shows association with chronic disease conditions. In amyotrophic lateral sclerosis (ALS) patients, researchers found a highly significant association between age acceleration and age at ALS onset, and greater age acceleration was associated with a shorter survival [457]. HIV patients show an increased Horvath age acceleration that is reversed following antiretroviral therapy [107, 353]. Curiously, female multiple sclerosis patients, compared to healthy females and all males, show a negative age acceleration by Horvath’s clock, although the authors also observe a change in blood cell composition in these patients that may be a contributory factor [397]. Among study participants with and without a mutation in the LRRK2 gene, Parkinson’s patients were found to be biologically older by Horvath’s clock, and that accelerated aging was further associated with an earlier age of onset, but not greater disease severity [396]. Non-alcoholic steatohepatitis patients with phase 3 fibrosis exhibit a higher epigenetic age, and age acceleration was significantly correlated with hepatic collagen content [243]. Roetker and colleagues report that increased carotid intima-medial thickness positively associated with Horvath age acceleration, while HDL cholesterol levels associated negatively [340]. They also observe increased hazards ratios for cardiovascular disease and related mortality in ten years with increased age acceleration. Age acceleration was significantly related to incident cardiovascular disease, with an increased risk of 4% of a cardiovascular disease event occurring with each year of increased biological age [235]. Individuals with increased pulse or increased systolic blood pressure, and women with hypertension were reported as having elevated epigenetic age in a 2022 study [445].

Conversely, a relatively large study involving the Rhineland Study population found no association with cardiovascular disease, and a separate study found no association with Horvath’s biological age was observed in ischemic stroke patients [119, 379]. A study of the Melbourne Collaborative Cohort Study population found an association with increased age acceleration and diabetes, but a much smaller study group found no such association [98, 419].

Physical parameters have been shown to associate with Horvath pan-tissue age acceleration, in particular BMI and related measures of physical mass [64, 110, 114, 140, 197, 226, 343, 370]. However, a twin study published in 2022, confirming that women were biologically younger than men according to Horvath’s clock, observed that that phenomenon was partially mediated through BMI, with men in the study population typically having higher BMI than women [181]. In women among the SAPALDIA and ECHRS cohorts, FEV1 and forced vital capacity (FVC) were associated with age acceleration, while in men there was an association with FVC in the follow-up time points only [335]. Related, levels of physical activity associate negatively with age acceleration in some, but not all, studies [114, 342, 381, 412, 419].

Among veterans diagnosed with COPD, it was observed that biological age was inversely associated with performance in the six minute walk distance test [425].

In a study involving bariatric surgery patients, a reduction in biological age was seen in patients following surgery [120]. In a study of overweight and obese breast cancer survivors weight loss, metformin treatment, or combination therapy did not result in an improvement in Horvath’s biological age, however [296]. Dietary improvement also resulted in deceleration of Horvath biological age: a randomized clinical dietary trial including probiotics and phytonutrients resulted in decreased biological age after eight weeks of intervention, and a trial involving adherence to a Mediterranean-style diet led to improvements in biological age among Polish participants, particularly women, after one year [115, 130]. However, other studies of the Mediterranean diet, the DASH diet, and the CALERIE trial found no significant changes in biological age [199, 431]. A study involving the Melbourne Collaborative Cohort found elevated meat consumption was associated with increased Horvath age acceleration [98]. Additionally, dietary supplementation has been found to influence biological age. A 2019 randomized clinical trial and a 2022 study both found vitamin D supplementation resulted in a decreased deltaAge among participants [63, 411]. Personal environment is relevant to biological age, as observed in a study of the Detroit Neighbourhood Health Study: neighbourhood quality was found to be associated with accelerated Horvath aging [260]. Two studies found a negative association between educational attainment and epigenetic age acceleration [98, 114]. Other work reported no association with educational attainment, childhood and adulthood socioeconomic status and changes in class, and household earnings [17, 131, 354]. Interestingly, one study identified an improvement in epigenetic age associated with having endured childhood financial hardship [110]. A 2022 report identified a deceleration of biological age among married compared to never married and divorced study participants [17]. No impact from spousal, child, friend, other family support or contact frequency was observed on Horvath biological age in a study examining the HRS population [157].

Alcohol consumption and substance abuse disorder diagnosis do not exhibit a correlation with Horvath’s biological age [42, 51, 198]. In several studies current or past smoking associates with increased biological age by the pan-tissue clock [110, 212, 419]. Similarly, airborne pollution is associated with increasing biological age acceleration: elevated PM_2_._5_, sulfate and ammonium exposure were associated with increased age acceleration [293]. The finding of PM_2_._5_ exposure leading to increased age acceleration was also found in a 2016 study that combined the NAS and KORA F4 study cohorts; this study also found that in men, increased PM_10_ exposure, while in women, black carbon and nitric oxides exposure were associated with increased age acceleration [428]. In a 2022 study, increases in PM2.5 exposure in the twenty four to forty eight hours prior to blood draw resulted in an increase in biological age [127]. Among the Lothian Birth Cohort of 1936, exposure to increased air pollution in young through middle adulthood was associated with an increase in Horvath’s biological age years later [18]. Pollution-related studies also show that while urine concentrations of arsenic, cadmium, lead, manganese and mercury did not influence age acceleration among the NAS population, blood levels of vanadium, cobalt, zinc and barium caused negative age acceleration, and nickel and arsenic levels were associated with positive age acceleration in a Chinese cohort [294, 443]. An American study found that among a population highly exposed to polybrominated biphenyl also exhibited accelerated aging according to Horvath’s pan-tissue clock [81].

Reproductive health has been found to influence biological aging. Studies of women have found positive associations with Horvath biological age acceleration and age at menarche, age at menopause, time since menopause or surgically induced menopause (bilateral oophorectomy) [64, 221]. However, no association between Horvath’s biological age and pre-eclampsia diagnosis was reported [319]. Epigenetic age association was found to increase slightly per live birth, and a woman’s age at first birth was inversely associated with age acceleration [196]. Gestational impacts to biological age can also persist throughout life: in a study of Canadian adults, men who had been born at an extremely low birth weight exhibited an advanced epigenetic age relative to men who were born at normal birth weight, or women regardless of birth weight [407]. An additional study found a similar result linking extremely low birth weight to later advanced epigenetic age, but a third paper published in 2022 found no association [204, 266].

Brain-related and mental health also exhibits associations with biological age. Data from the San Antonio Family Study found that global white matter index was negatively correlated with age acceleration. An MRI study of 79 participants found that cortical thickness decreased with advancing combined Horvath/Hannum biological age [318]. A study of an extended Mexican-American pedigree found an association with reductions in white matter integrity in specific brain regions and accelerated Horvath age, with evidence that this is due to underlying genetic influences [159]. However, the findings associating cognitive capacity with changes in biological age using Horvath’s pan-tissue clock have been mixed. Work from the HANDLS and Middle-Aged Danish Twin Study cohorts found no association with cognitive function and Horvath biological age [36, 383]. Conversely, a relatively small study of 68 cognitively healthy adults found that selective attention performance was better predicted by a combined Horvath/Hannum biological age than by chronological age [435]. The VITAL-DEP study found an inverse association with objective cognitive score and accelerated biological aging [419]. In their study, Cruz-Almeida and colleagues found lower fluid cognition associated with greater biological age [80]. They also observed that greater emotional stability, conscientiousness, and lower extraversion associated with a younger epigenetic age, while a lower heat pain threshold and pressure pain threshold at the trapezius was associated with a greater epigenetic age. However, a number of studies report no association between Horvath’s biological age and cognitive decline or decline in mental facility [303, 359, 390, 412]. Conflicting results were found in studies of schizophrenic individuals: a 2020 study found that male and female schizophrenics exhibited increased age acceleration, and that males taking clozapine alone or in conjunction with antipsychotic medication partially ameliorated that acceleration, while a 2018 study had found that in post-mortem brain samples there was no difference in biological age relative to controls, while blood samples from male schizophrenics exhibited reduced age acceleration, with no corresponding effect seen in female schizophrenics [156, 274]. A 2021 study found no significant difference between controls and schizophrenic patients in terms of biological age, but did report Horvath age acceleration correlating with psychotic score [82]. The absence of a significant association between Horvath’s biological age and hypersexual disorder was reported in a 2022 paper [43].

No associations between autism diagnosis or ADHD genetic burden have been reported [10, 301]. While two studies observed an association with Horvath age acceleration and post-traumatic stress disorder, a study of caregivers of autistic children found no increase in biological age associated with the stress of caring for a child with autism as opposed to a neurotypical child [143, 339, 427]. A 2019 study found an association with increased age acceleration and anxiety, but not with diagnosis of depression [419]. Mental health interventions appear to be beneficial: a study of meditators found that each year of regular meditative practice was associated with a decrease in age acceleration among older, but not younger, meditators, and a randomized controlled trial including relaxation activities resulted in younger epigenetic ages relative to controls following 8 weeks of practice [62, 115]. Lastly, studies on sleep quality have revealed mixed results. A study involving participants from the WHI found no impact of sleep disturbance on age acceleration, while a study combining data from the MESA and Framingham Heart Study (FHS) populations found that most disordered breathing traits were associated with epigenetic age acceleration in men, but not women, although the associations of sex and sleep disturbances were not as clear with further statistical analysis [55, 227].

#### Horvath’s Skin and Blood Clock

In contrast to the widespread use of Horvath’s pan-tissue clock, Horvath’s skin and blood clock, developed in 2018 using 391 CpGs, has yet to achieve similar levels of adoption [167]. In preparation for this review, we identified eleven papers that make use of the skin and blood clock. Treatment of skin biopsies with rapamycin, a senomorphic agent, revealed a biological age increase according to the skin and blood clock, contrary to the finding by a newly described skin-specific epigenetic clock [41]. In a tissue culture study, the Horvath skin and blood clock was found to report the equivalent of 25 years of aging over the 152 day long course of growth of cultured fibroblasts, including during senescence [389]. The study also found that fibroblasts grown under hyperglycemic conditions exhibited the equivalent of 3.4 years elevation in predicted biological age.

However, a study of type I diabetics found that while the skin and blood clock did correlate with chronological aging, it predicted lower than chronological age in those participants, and no association with risk of incident cardiovascular disease, diabetic retinopathy, or neuropathy [342]. Lower skin and blood clock age was associated with increased levels of physical activity, and higher age was associated with increasing BMI. In a study of the HRS population, white study participants were found to be biologically older than other participants, while for non-white participants, there was an association of experience of discrimination with elevated epigenetic age [37]. No associations between the skin and blood clock were identified for substance abuse disorder, early ovarian aging, weight loss and/or metformin therapies, autism diagnosis, pre-eclampsia, socioeconomic status, and survival of head and neck cancers [51, 69, 296, 301, 319, 354, 444].

#### Hannum’s Clock

Hannum’s epigenetic clock was the first multi-CpG clock published, in early 2013, and relies on the methylation signatures of 71 CpG sites [147]. The clock was developed using blood methylation data exclusively, rather than employing a multi-tissue approach to estimating biological age. Of the 75 papers reviewed here that make use of the Hannum clock, all of them that reported it showed a positive correlation with chronological age; further, all that reported a correlation found a positive association with the biological age predicted by Horvath’s pan-tissue clock as well [30, 64, 82, 98, 225, 256, 340, 379]. Hannum epigenetic age was not found to associate with facial aging [257]. As was observed with the Horvath pan-tissue clock, there have been sex-specific differences reported, with men exhibiting a higher Hannum age than women [17, 18, 37, 98, 110, 130, 204, 228, 260, 390, 411]. Also as observed with the Horvath pan-tissue clock, white study participants exhibited a higher Hannum age acceleration than did African American study participants [394]. Hannum clock-measured biological age has an estimate of heritability of 42%, and is found to be significantly correlated in monozygotic twin pairs as compared to non-twin sibling pairs [225, 256]. In a study of middle-aged Danish twins, the Hannum measure was found to overestimate the age of study participants more often than underestimate age [383]. The Hannum clock correlates strongly when measured on the Illumina Infinium Human Methylation 450k BeadChip and the Infinium MethylationEPIC BeadChip [272]. Sturm and colleagues found that Hannum’s clock predicted a linear age increase in cultured fibroblasts up until the senescence phase, when the clock ceased to progress [389].

Studies have found elevated Hannum age or Hannum age acceleration associated with increased risk of all-cause mortality [110, 228, 256]. Among patients with head and neck cancer, increased morbidity was associated with elevated Hannum age in a 2021 study [444]. In studies of ischemic stroke survivors, Hannum age acceleration was observed in survivors as compared to controls, with stroke survivors exhibiting an age 2.5 years older as measured by the Hannum clock [380]. Elevated Hannum biological age, but not Horvath pan-tissue age, was associated with poorer outcomes three months following stroke [380]. Increased levels of markers of cardiovascular dysregulation were associated with higher Hannum age acceleration in women only [269]. Increased risk for cardiovascular disease and mortality in the first and second ten years following initial observation were all associated with Hannum age acceleration in a study of middle-aged African Americans [340]. However, in a study of Swedish senior citizens, no relationship between Hannum age and cardiovascular disease was detected [235]. In unadjusted models, Roberts and colleagues report a link between Hannum age acceleration and atrial fibrillation in a 2021 study [337]. Decline in FEV1, a marker of pulmonary health, was associated with Hannum age among female but not male participants of the SAPALDIA and European Community Respiratory Health Survey (ECRHS) cohorts [335]. In a study of twins, the twin exhibiting higher Hannum age acceleration was more than twice as likely to die first than the lower Hannum aged twin [70]. Work with the Irish Longitudinal Study on Aging population found an association between increased Hannum age acceleration and the likelihood of taking five or more medications daily, a proxy feature of multimorbidity [270]. Members of the Lothian Birth Cohort of 1936 were significantly more likely to be frail when they exhibited Hannum age acceleration [122, 123]. In a study of postmenopausal women, it was found that the odds of surviving to age 90 with intact mobility worsened with increased biological age [171]. Conversely, other studies have found no association with Hannum biological age and impairment of activities of daily life or physical decline [139, 412]. Belsky and colleagues report Hannum age acceleration associated with most of the markers of decreased health span measured in their study of 1037 members of the Dunedin Study cohort [30]. In terms of physical activity, results are mixed: a study of veterans with COPD revealed an improvement in Hannum biological age with an increase in exercise capacity, but other studies report no association with levels of physical activity [119, 381, 425].

As observed with the Horvath pan-tissue clock, a number of physiological features associate with changes in Hannum biological age. Chiefly among them is BMI and related features including waist to hip ratio and waist circumference [98, 114, 197].

Diabetes and gestational diabetes diagnoses were also associated with Hannum age acceleration [196, 340]. Elevated HbA1C was associated with increased Hannum age acceleration, particularly in men, and metabolic dysregulation biomarkers were associated with increased Hannum biological age in both sexes [269]. Features of reproductive health associate with the Hannum clock: a woman’s age at first live birth was associated with decreased Hannum age acceleration, while a small age acceleration was associated with each additional live birth [64, 196]. There was no association observed with diagnosis of pre-eclampsia during pregnancy, nor with maternal energy reserves as represented by arm fat [204, 319]. Further, there was no association detected between Hannum’s biological age later in life and birth weight [204].

Work with the Melbourne Collaborative Cohort found that increased dietary fruit intake was associated with decreased Hannum age acceleration, but other studies analyzing the effects of adherence to a Mediterranean diet did not improve Hannum age [98, 130, 199]. Weight loss regimens, including the CALERIE intervention program and a combination therapy including metformin and weight loss, did not impact Hannum epigenetic age acceleration [296, 431]. Dietary supplementation of vitamin D was also found to be associated with improved Hannum age, although serum levels of 25-hydroxyvitamin D were not; a later study did not replicate the finding linking vitamin D levels to epigenetic age [63, 411]. Beach and colleagues found in a study of the Family and Community Health Studies (FACHS) cohort that 55 of the 71 CpGs of the Hannum clock associate significantly with alcohol consumption, while 33 of the 71 CpG sites respond significantly to smoking; while smoking always led to Hannum age acceleration, moderate levels of alcohol consumption reduced Hannum age, while very low and very high levels of alcohol consumption were accelerative [23]. A later study of women participating in the Sister Study reaffirmed that current alcohol consumption was associated with increased Hannum epigenetic age [198]. Several other studies have noted the association between current or prior smoking and Hannum age acceleration [98, 114, 212, 340]. In a small study of 22 current smokers, successful smoking cessation led to a decrease in Hannum age acceleration [212]. A study of Finnish mixed-gender twin pairs indicated that, while the male twin was biologically older, this effect was largely mediated by an increased likelihood to be a smoker relative to his sister [181]. In terms of air pollution and its impact on biological age, results are mixed: one study found no association with historical air pollution levels and Hannum age acceleration later in life, while a separate study identified a detrimental impact of immediate exposure to air pollution prior to testing [18, 127]. A study of a population exposed to polybrominated biphenyl found that Hannum age acceleration associated with increasing exposure [81]. Similarly, exposure to certain organochlorine pesticides were associated with accelerated Hannum age in Swedish senior citizens [236]. A study of long-term residents of Guangxi, China found that blood serum levels of vanadium, cobalt, zinc and barium associated with a decrease in Hannum age acceleration, while levels of nickel and arsenic accelerated Hannum age [443].

Studies of mental health have revealed associations with the Hannum epigenetic clock. In a comparison of schizophrenic and control individuals, Hannum age acceleration was observed in schizophrenics over controls [82]. Male, but not female, schizophrenia patients taking clozapine or clozapine in conjunction with an antipsychotic medication showed a decrease in Hannum biological age relative to those not taking clozapine [156]. Anxiety, but not depression, was associated with Hannum age acceleration in a large 2020 study, but a later study of twins revealed a higher Beck Depression Inventory II score associated with elevated Hannum biological age [240, 339]. In men, accelerated Hannum aging was associated with a faster decline in memory, visuoconstructive ability, attention and processing speed, but a previous study in a larger population found that Hannum biological age was not a good indicator of cognitive change in the middle aged [36, 383]. Additional studies revealed no apparent association between cognitive performance and Hannum’s clock [303, 359, 412]. However, among people living with HIV, decreasing neural performance was associated with elevated biological age [353]. Autism, hypersexual disorder, and substance abuse disorder showed no association with Hannum’s clock, but a twin study revealed that twins with PTSD were biologically older than their unaffected sibling [43, 51, 301, 427].

Low income and financial pressure have both been associated with accelerated biological aging [213, 339, 371]. Similarly, neighbourhood quality has been associated with Hannum age acceleration [213, 260]. In terms of socioeconomic status, results have been mixed: some studies report no significance, while others report that reduced childhood social class or economic status is associated with accelerated Hannum age, although one 2023 study reports an inverse finding for childhood financial hardship resulting in decelerated aging [17, 110, 131, 354]. A study of the HRS population reported the absence of an association with the experience of discrimination, and a later study of the same population detected an increase in familial contact associated with a lower Hannum age [37, 157]. Finally, Hannum biological age has been shown to associate negatively with educational attainment [114, 340].

#### PhenoAge

In 2018 Levine and colleagues published their second generation epigenetic clock based on 513 CpGs, referred to as PhenoAge [223]. This clock was developed using DNA methylation data from whole blood, and is designed to collect methylation status not only from chronological age-linked CpG sites, but also those sites that are responsive to changes in physiological status of an individual. Nonetheless, PhenoAge does correlate well with chronological age in blood and other cell types, and among the 70 papers reviewed for their use of PhenoAge, all that reported it found an association with chronological age. Furthermore, PhenoAge shows correlation to varying degrees with first generation clocks such as Horvath’s pan-tissue and Hannum’s [51, 64, 95, 143, 294, 342, 427]. PhenoAge also shows correlation with other second generation clocks, such as DunedinPoAm and GrimAge [31, 51, 95, 294, 342, 427]. Clocks not extensively based on DNA methylation, such as measures of allostatic load and metabolomic metrics, show association with PhenoAge [269, 339]. It also correlates negatively with mtDNA copy number, but studies considering telomere length and PhenoAge are mixed [51, 95, 149, 357]. Contrary to findings in other clocks, two studies report finding lower PhenoAge in men relative to women; Li and colleagues found that while increased PhenoAge was associated with increased mortality risk, the effect was much stronger in women than in men [204, 228, 260]. However the majority of studies that report a sex-based difference find women with a younger PhenoAge than men [12, 18, 110, 181, 303, 390, 411]. The results of ethnicity-based analyses are mixed, and limited: five studies report ethnicity-based PhenoAge, with one finding Hispanics age at a slower rate than non-Hispanics, two reporting no association with race or ethnicity among study participants, and two reporting higher PhenoAge acceleration among Black participants as compared to Whites or Hispanics [12, 37, 110, 139, 200]. A study of cultured fibroblasts found that PhenoAge increased linearly over the growth phase of the cells, but ceased to increase once the cells reached senescence [389]. Two of the studies reviewed here found that PhenoAge systematically underestimates chronological age, one examining biological age among HIV patients prior to and during antiretroviral therapy, and one examining a large cohort of older adults [107, 368]. However, PhenoAge does associate with markers of multimorbidity, limitations in activities of daily life, cognitive function, and increased risk of mortality [78, 125, 139, 149, 228]. A study of postmenopausal women associated elevated PhenoAge acceleration with a decrease in the likelihood of surviving to age 90 with intact mobility and cognitive function [171]. In contrast, a study of 490 participants in the Irish Longitudinal Study of Aging found no association between PhenoAge and any of the physical or cognitive outcomes measured in that study, and a study of the BASE II population had similar non-significant findings [270, 412]. Both poorer initial performance and accelerated decline in memory function tests are associated with worsening PhenoAge acceleration, although the effect is partially mediated by socioeconomic status [12]. Additional studies considered for this review find no association with cognitive functioning or dementia with PhenoAge acceleration [303, 359, 390].

Individual physical parameters associate with changes in PhenoAge. Sleep quality in terms of a higher apnea/hypopnea index and arousal index associated with PhenoAge acceleration in women but not in men in the MESA population, but the inverse sex association was found in the FHS population [227]. In the UK Biobank population, an early chronotype, and achieving normal sleep duration were associated with a decelerated PhenoAge [126]. That study also observed that daytime sleepiness and difficulty rising was associated with an accelerated PhenoAge. Serum levels of vitamin B6 negatively associate, while serum folate levels positively associate with PhenoAge; vitamin D supplementation did not influence PhenoAge in either healthy or vitamin D deficient study participants in a separate study [295, 411]. As with other clocks described here, BMI associated positively with increasing PhenoAge, as do waist to hip ratio and waist circumference [53, 64, 110, 114, 181, 197, 200, 444]. Changes in BMI from non-obese to obese resulted in age acceleration in a study of the NHANES population [53]. In comparing men and women, it was found in a Finnish twin study that the tendency for males to exhibit a higher PhenoAge than females is partially modulated by an associated higher BMI observed in male study participants [181]. An intervention trial comparing conventional weight loss with metformin treatment or combination therapy found no improvement in PhenoAge acceleration among overweight and obese female study participants; similarly, dietary intervention in the CALERIE trial did not affect PhenoAge [296, 431]. Conversely, the Sister Study population found that adherence to a Mediterranean or DASH diet was associated with lower PhenoAge, particularly among women with low physical activity [199]. Higher levels of daily physical activity, including increased six minute walk distance and increases in grip strength, were reported to associate with decreased PhenoAge [119, 241, 357]. In a longitudinal study of pregnant women, it was found that increases in BMI during pregnancy were associated with lower post-birth PhenoAge, but that having no decrease in BMI in the year post-birth was associated with a higher post-birth PhenoAge [343]. Diagnosis of pre-eclampsia resulted in an elevated PhenoAge as compared to normotensive pregnant women [319]. Self-reported abnormal glucose tests during pregnancy and diagnosis of gestational diabetes were found to be associated with increased PhenoAge, but in a cell culture study, growth in hyperglycemic conditions did not lead to a PhenoAge acceleration [196, 389]. Among women with early ovarian aging, no difference in PhenoAge was detected relative to women with normal ovarian aging [69]. Additionally, other features of reproductive health associate with PhenoAge: PhenoAge acceleration occurs slightly after each live birth, but was inversely correlated with a woman’s age at first live birth [196]. There was no observed association with age at menarche or other features of pubertal timing [143]. Among males only, birth weight was inversely associated with PhenoAge later in life [204].

Chronic disease is associated with a higher PhenoAge. Multiple sclerosis patients were found to have a higher PhenoAge independent of their BMI or smoking status [397]. Studies of HIV positive individuals show that low CD4+ cell counts show significant PhenoAge acceleration, and antiretroviral therapy leads to a decrease in PhenoAge [107]. In schizophrenia, a higher PhenoAge is observed in patients as compared to controls, but unlike the results found for the Horvath pan-tissue and Hannum clocks, no decrease in PhenoAge was observed among patients on clozapine or clozapine in conjunction with antipsychotic medications [156]. No association was observed with diagnosis of autism or hypersexual disorder, but a higher Beck Depression Inventory II score was associated with an accelerated PhenoAge [43, 240, 301]. Among individuals with substance abuse disorder, an elevated PhenoAge was detected relative to controls in blood samples, but not in post-mortem brain samples [51]. A twin study revealed that twins with post-traumatic stress disorder (PTSD) have an elevated PhenoAge as compared to their healthy twin [427]. Roshandel and colleagues found in a study of type I diabetics that repeated albumin excretion rate, a marker of kidney function, was associated with increased PhenoAge, but other diabetic complications such as diabetic peripheral neuropathy and cardiovascular autonomic neuropathy were not [342]. Metabolic dysregulation, a subtype of dysregulation under the allostatic load measurement, was found to be correlated with increased PhenoAge in both men and women, while cardiovascular dysregulation was only significant in women [269].

Hypertension and atrial fibrillation been observed in association with increased PhenoAge acceleration [337, 339]. In a relatively small study, chemotherapy treatment was not found to associate with increased PhenoAge acceleration, while for radiation therapy, increased age acceleration at 6 and 12 months post-therapy were associated with worse overall survival and progression free survival [241, 444].

Environmental and lifestyle exposures also influence PhenoAge, in particular smoking and heavy alcohol consumption [42, 64, 110, 114, 200, 339, 374, 375]. Work on the Sister Study population, an all-female data set, did not confirm an association with alcohol consumption and accelerated PhenoAge, however [198]. A Finnish twin study found that while male twins were more likely to have a higher PhenoAge than their sisters, this affect was modulated by unhealthy lifestyle habits including smoking and alcohol use [181]. Study of those exposed to polybrominated biphenyl found PhenoAge increased with increased exposure [81]. Urinary metal concentrations measured in participants of the NAS was found to associate with PhenoAge acceleration, in particular levels of manganese alone, or in a mixture model a combination of arsenic, cadmium, lead, manganese and mercury [294]. Air pollution was not found as significant in a study of the Lothian Birth Cohort of 1936, but work with the UK Biobank population did find an association with higher PM2.5 and atmospheric NO2 levels and increased PhenoAge [18, 126]. There was no impact detected from immediate exposure within the last forty-eight hours prior to testing in a population of Chinese students, however [127]. Beyond exposure to pollution, neighbourhood quality was found to have a significant association with PhenoAge, although only in women; a later study had similar findings, albeit with no gender effect [214, 260]. Income and educational attainment are both negatively associated with PhenoAge [17, 114, 131, 200, 339]. Other measures of socioeconomic status, both during childhood and adulthood, associated with PhenoAge [17, 110, 131, 354]. Other childhood experiences, including childhood adverse experiences, have been studied in relation to accelerated PhenoAge later in life, although the findings have been limited: one study detected an association with abusive, but not neglectful, childhood maltreatment associated with accelerated biological aging, but other studies report no association with adverse experiences or childhood police encounters [37, 85, 176, 325]. No association was found between perceived discrimination or social isolation and elevated PhenoAge [37, 85, 157].

#### GrimAge

GrimAge is a recently developed second generation epigenetic clock that includes CpG methylation estimates of seven key physiological risk and stress-related plasma proteins, as well as a CpG-based indicator of total smoking pack years [245]. In total it assesses methylation at 1030 CpG sites and is especially accurate at estimating time to mortality. A relatively recently developed clock, 65 papers studied in this review employed GrimAge as an epigenetic age metric, and for the papers that reported it, GrimAge was found to correlate with chronological age. Three studies reported a significant correlation in predictions of GrimAge with both Horvath’s pan-tissue and PhenoAge [294, 296, 342]. An additional study tested the relationship between mtDNA copy number and GrimAge and found no association [95]. One study reviewed here found that GrimAge tended to overestimate age [107]. Elevated GrimAge was found in men relative to women, and lower in white individuals as compared to other ethnicities [10, 12, 18, 139, 143, 181, 200, 204, 275, 303, 304, 316, 373, 375, 390, 411]. GrimAge and GrimAge acceleration strongly associate with increased risk of mortality: two studies associated a 39% and 91% increase in mortality associated with one standard deviation increase in GrimAge, the former study observing a stronger effect in women than in men [228, 270]. Accelerated GrimAge was associated with decreased facility in the activities of daily life, with poorer healthspan-related measures and with a decreased probability of achieving age 90 with intact motility [110, 139, 171]. Conversely, a 2022 study did not find GrimAge as a good marker of either physical or mental health deterioriation [412]. Increased GrimAge is associated with more diagnoses of chronic disease as well as with taking five or more medications daily, a proxy measure for chronic disease [270, 275]. In particular, development of diabetes in patients with obesity and/or hyperglycemia, as well as the incidence of three key diabetic complications, were associated with elevated GrimAge acceleration [190, 342]. Increased likelihood of incidental atrial fibrillation was found to be associated with elevated GrimAge in a 2021 study [337]. Studies of lung health in a large, combined cohort found an association of GrimAge with reduced FEV1, FVC, as well as the ratio of FEV1/FVC [335]. There was no association of GrimAge and pre-eclampsia compared to normotensive pregnancy identified in a 2022 study [319]. Studies on GrimAge in other chronic diseases are limited. Higgens-Chen and colleagues found GrimAge acceleration in schizophrenic patients versus controls [156]. Increased ADHD polygenic score is associated with an accelerated GrimAge; in a study of high functioning autistics, only one component of GrimAge (DNAmPAI-1) was significantly associated, with GrimAge overall having insignificant associations [10, 301]. Patients diagnosed with hypersexual disorder did not exhibit elevated GrimAge relative to healthy controls [43]. In a study of HIV positive individuals, significant GrimAge acceleration was observed among those with a high viral load, which improved following antiretroviral therapy [107].

In terms of cognitive health, a number of studies have investigated various facets of mental and neurological health. Findings related to depression are mixed: a 2023 report identifies an association with increased GrimAge that is not supported by an earlier report finding no connection between depression severity and GrimAge [110, 240]. Three studies investigating trauma-exposed and PTSD-diagnosed veterans found differing patterns of association as well: a 2021 study affirmed a link between GrimAge acceleration and PTSD diagnosis and severity, while a 2022 study found the PTSD link only significant among younger veterans, and a 2023 study found a link with PTSD but no improvement in GrimAge age acceleration following treatment [153, 184, 447]. The 2022 study also observed that younger veterans had significant association between cognitive disinhibition and poorer memory with GrimAge, while inflammatory and neuropathological traits were associated regardless of chronological age. When comparing nonpathological differences in cognitive ability, it was found that women possibly exhibited a younger GrimAge than men due to differences in those abilities [303]. A further study of GrimAge in American veterans uncovered links with educational attainment, marital status, combat history, accumulated past traumatic experiences and substance abuse disorder with worsening GrimAge, while increases in physical activity frequency, improvements in sleep quality, and a predisposition towards openness and gratitude were all found to be protective [395]. In a study of APOE *ε*4 status, it was found that there was an interaction with verbal memory among women and GrimAge acceleration that was not observed in men [304]. Poorer initial memory performance and a steeper decline in performance are both associated with accelerated GrimAge, a finding that was partially mediated by socioeconomic status [12]. There have been no studies positively identifying an association with GrimAge acceleration and diagnosis of either mild cognitive impairment (MCI) or dementia, however [359, 390].

Studies concerning the relationship between BMI and GrimAge are mixed. In a study of 2758 non-Hispanic, middle-aged white women, GrimAge was found to be associated with BMI, waist to hip ratio, and waist circumference, while a smaller study found no association with BMI [197, 270]. Later studies are more decisive, finding a positive association between BMI and accelerated GrimAge [110, 182, 447]. Dietary interventions have also been mixed: a 2023 study of the CALERIE intervention found no significant effect, while a 2022 investigation of the Sister Study population found both the DASH and Mediterranean diets associated with decreased GrimAge [199, 431]. A trial involving metformin, conventional weight loss, or combination therapy in overweight and obese breast cancer survivors did not find an association with GrimAge, however [296]. In a study of pregnant women, it was found that there was a significant decrease in GrimAge measured at post-birth relative to during pregnancy, that increases in BMI during pregnancy were associated with a lower post-birth GrimAge, but that having no decrease in BMI in the year post-birth was associated with a higher GrimAge [343]. Birth weight is not reported to be associated with GrimAge [204]. In terms of blood-based metabolites, Nwanaji-Enwerem and colleagues found that levels of vitamin B6 are associated with decreased GrimAge, while folate levels are associated with elevated GrimAge [295]. In a study of the BASE II population, vitamin D levels were found to be inversely associated with GrimAge [411]. History of and current smoking, as well as elevated alcohol consumption, were associated with increased GrimAge [42, 110, 181, 184, 198, 228, 275, 373–375, 447]. However an association between GrimAge and substance abuse disorder in either blood or brain samples was not detected [51]. Physical activity and elevated leisure activity were found to be associated with lower GrimAge acceleration among men in a Finnish twin study, but a more focussed study comparing master rowers to age-matched sedentary controls did not find an effect of physical activity or prowess [181, 357]. Other studies found the influence of increased physical activity on GrimAge deceleration, including higher daily step counts, improvements and ability in the six minute walk distance test, and associations with hand grip strength were all reported [119, 241, 316, 381, 425]. Joint pain, specifically low and high impact knee pain, conferred an older GrimAge biological age, and this partly mediated the effect of pain on balance performance [315].

Socioeconomic features have been studied in conjunction with GrimAge. McLachlan and colleagues found elevated GrimAge with lower levels of educational attainment, lower social status, and psychosocial loss [275]. Simons and colleagues found associations with educational attainment, household income, neighbourhood disadvantage and cumulative disadvantage [373]. Neighbourhood and historic pollution levels have had mixed findings: a study of the Lothian Birth Cohort of 1936 found no influence on GrimAge of historical pollution levels, while a study of Chinese students found that an increase in PM_2_._5_ immediately prior, or in the window of time 24 to 48 hours prior to examination, increases GrimAge [18, 127]. Higher levels of early life abuse and trauma were associated with elevated GrimAge acceleration later in life [143]. An additional study identified an association between cumulative adverse childhood experience score and GrimAge, but this finding was not corroborated by an earlier 2022 study [176, 325]. It was also found parental separation or divorce and childhood emotional abuse also correlated with accelerated GrimAge later in life [176]. Overall, both childhood and adult socioeconomic status associated with GrimAge, although the effects were found to be additive rather than multiplicative [131, 354]. Lower educational attainment and neighbourhood disadvantage are both associated with accelerated GrimAge [35, 131, 200, 214]. GrimAge is significantly associated with both childhood police encounters and previous incarceration [34, 35, 85]. Mental health considerations also factor into GrimAge and GrimAge acceleration: more symptoms of depression, as well as a negative attitude towards physical changes associated with aging, are linked to GrimAge acceleration [275]. Perceived everyday discrimination and cumulative disadvantage also associate with elevated GrimAge [37, 374, 375]. Studies of African American youth found that changes in self control ability during late adolescence was associated with an improved GrimAge in early adulthood [215]. Higher levels of support or contact with friends, or higher contact with one’s children, was associated with decelerated GrimAge [157].

#### Weidner’s Clock

In 2014, following the Hannum and Horvath pan-tissue epigenetic clocks, the Weidner clock was developed. This clock was originally developed with the intention to interrogate CpG methylation status via bisulphite pyrosequencing, rather than the then-costly microarray approach [433]. The Weidner clock is derived from the methylation status of 102 CpG sites, 99 of which are present on the Illumina Infinium Human Methylation 450k BeadChip; many of the reviewed uses of the Weidner clock rely on microarrays and those 99 CpG sites rather than the complete 102 CpG clock.

Weidner and colleagues further distilled their clock down to a 3 CpG clock that was nearly as predictive as the complete 102 or 99 CpG sets; those three CpGs were associated with the genes PDE4C, ITGA2B and ASPA. Six papers reviewed here make use of the Weidner clock, and the three that assessed association with chronological age found a positive association [116, 230, 271]. One paper reported a weak correlation between the Weidner 99 CpG set and biological age as measured using 10 biomarkers and the Klemera-Doubal method, but none of the studies considered here reported any correlation with any of the other methylation-based clocks [30]. A study of the combined Lothian Birth Cohorts of 1921 and 1936 found that per five year age acceleration according to the 99 CpG clock, there was an associated 11% increase in risk to mortality, however, only one of the 3 CpG clock sites, the CpG close to the gene PDE4C, was significantly associated with survival [230]. In the Lothian Birth Cohort of 1921 alone, there was no association between Weidner epigenetic age and estimated face age, and a study of the Dunedin cohort found no association with any of the healthspan-related characteristics they measured [30, 257]. No correlation was found with either the 99 or 3 CpG versions of the clock with experience of discrimination in a study of the HRS population [37].

#### Pace of Aging and DunedinPoAm

The ‘Pace of Aging’ epigenetic clock, which measures the rate of physiological deterioration, was developed in 2015 by Belsky and colleagues, and is based on 18 biomarkers covering a variety of different organ systems, calculated using the Klemera-Doubal algorithm [27]. In that seminal paper, the authors compared their Pace of Aging clock against a 10 biomarker clock also calculated using the Klemera-Doubal method, and found they correlated positively and to a moderate degree. This paper also found that the Pace of Aging measure correlated negatively with a number of physical ability measures, and positively with an increased degree of physical limitation. Pace of Aging also correlated negatively with cognitive measures including cognitive function and cognitive decline. Lastly, the authors found that Pace of Aging associated with qualitative assessments of aging, including self-rated health and facial aging. In a follow-up study, Belsky and colleagues found that Pace of Aging was significantly correlated with the age of one’s longest-lived grandparent, childhood social class, health, IQ, self control and adverse childhood experiences [28]. This study was followed by a third, that found weak correlation between Pace of Aging and methylation-based epigenetic clocks and telomere length [30]. They also confirmed their earlier findings, with Pace of Aging significantly associated with physical functioning, cognitive functioning, and cognitive decline, as well as self-rated health and facial aging. A study of Finnish twins re-iterated the poor correlation with first generation epigenetic clocks and mild correlation with second generation clocks with the biomarker-based Pace of Aging [182]. A 2021 paper found that soluble urokinase plasminogen activator receptor (suPAR), a biomarker of inflammation and biological aging, was positively correlated with Pace of Aging [328]. Adolescent behaviors, including smoking, antisocial behavior, and obesity have been associated with accelerated Pace of Aging later in life [46, 205]. Socioeconomic disadvantage was also associated with accelerated biological aging, partly due to with accelerated aging, and this effect was partly modified by differences in health behavior correlated with socioceconomic status [355]. A quartet of unhealthy behaviors (obesity, low physical activity, elevated smoking and alcohol consumption) were also correlated with elevated Pace of Aging [182]. However, the presence of macular drusen, a retinal defect associated with elevated oxidative stress, were not associated with a difference in Pace of Aging biological age [437]. Using the Pace of Aging metric and facial aging measurements, work done in the Dunedin Study population found that participants with high quality relationships with low or absent levels of intimate partner violence had the lowest pace of aging compared to those in lower quality, higher violence relationships [45].

In 2020, Belsky turned his attention to methylation markers of biological aging, and using Pace of Aging as the criterion, used elastic net regression to identify 46 CpG methylation sites associated with the Pace of Aging metric [31]. Interestingly, none of the 46 CpG sites selected overlap with those found in the Horvath pan-tissue, Hannum, or Levine PhenoAge clocks. This novel clock was called DunedinPoAm for the study group used to develop it. It was found among Dunedin participants at age 45, relative to their measurements at age 38, that DunedinPoAm increases were associated with declines in physical and cognitive capability, and increases in physical limitations, facial aging and self-reporting of fair or poor health. Since its publication, usage of the DunedinPoAm clock has been steady. Sex-based difference in DunedinPoAm, where reported, has given mixed results [12, 17, 110, 204, 390]. Three studies report elevated DunedinPoAm epigenetic age in Black study participants than in other racial groups [12, 110, 200, 362]. A twin study found weak correlation of DunedinPoAm with first generation clocks and only modest correlation with second generation epigenetic clocks [182].

Unhealthy lifestyle factors including obesity, smoking, alcohol consumption and low physical activity are all associated with elevated DunedinPoAm [24, 110, 182, 200, 374, 375]. Conversely, dietary interventions such as the CALERIE trial resulted in improved biological age [431]. Birth weight was found to be inversely associated with DunedinPoAm age acceleration later in life [204]. Other physical parameters, such as post-intervention improvements in the six minute walk test distance and handgrip strength, were also associated with a decelerated DunedinPoAm [241]. Cognitive dysfunction is found to associate with accelerated aging: a 2022 study reported that dementia patients exhibited a higher biological age than did those with mild cognitive impairment, who in turn were biologically older than cognitively normal study participants [390]. That study also observed that the risk of dementia later in life as well as worse cognitive testing scores independent of diagnosis were both linked to elevated biological age. A study of the HRS population found that memory performance and rate of decline were associated with biological age, although the effect was partially mediated by socioeconomic status [12]. Applying DunedinPoAm to the NAS population found an association with chronic disease morbidity and increased risk of mortality [31]. A 2022 study reported an association between elevated DunedinPoAm and functional impairment, mortality, activities of daily life limitations, chronic conditions, and self-rated health [139].

Associations between elevated biological age and elevated levels of childhood deprivation and victimization were found among the E-Risk study population [31]. Other studies affirmed the link between childhood socioeconomic status and DunedinPoAm aging later in life, as well as other childhood incidents including police encounters and general childhood instability [17, 85, 110, 375]. Educational attainment is inversely associated with biological age [17, 110, 200, 354]. In adulthood, socioeconomic status and household income and wealth, including changes in income, continue to associate with DunedinPoAm [24, 354, 362, 375]. While cumulative disadvantage did associate, the experience of discrimination was not found to be associated with biological age [37, 354, 374]. Divorce was found to accelerate DunedinPoAm [17].

Neighbourhood features, including neighbourhood disadvantage, are also linked to changes in biological age [214, 354]. An increase in pollution exposure in the 24 to 48 hours immediately prior to testing also resulted in an elevated DunedinPoAm biological age [127]. Lastly, personal experience parameters, such as reductions in perceived loneliness and higher support and contact with friends, were related to lower levels of biological aging [24, 157].

#### Klemera-Doubal Algorithm

While not strictly a biological age clock, the Klemera-Doubal method (KDM) or Klemera-Doubal algorithm is often employed in conjunction with varying clinical biomarkers to assess biological age [192]. While a complete characterization of KDM is beyond the scope of this review, two key features should be highlighted. Klemera and Doubal conceptualized the problem as one of minimizing the distance between regression lines and biomarker points within the multidimensional space of all analyzed biomarkers, and they recognized that chronological age itself could be recognized as a biomarker to measure biological age, although this last point is not without controversy. KDM has been employed steadily since its initial publication with a variety of overlapping sets of biomarkers. Originally proposed as an improvement upon multiple linear regression (MLR) based approaches, this premise has since been tested and found valid [173, 219, 467]. In a 2013 publication, Levine compares KDM, MLR and principal component analysis (PCA) and found that a set of 7 biomarkers selected by PCA and used in a KDM calculation of biological age performed best in terms of mortality prediction [219]. A later paper compared the three approaches with a more expansive set of 68 clinical biomarkers and found that KDM-based biological age was the best predictor of frailty, especially in men; for mortality, all three algorithms outperformed chronological age [467]. In a study of 3642 women, Jee and colleagues similarly found that in a comparison of KDM, MLR and PCA, KDM performed as well as PCA [173].

In studies comparing KDM-calculated biological age to other epigenetic clocks, findings are generally of a modest or weak correlation between Horvath’s pan-tissue clock, Hannum’s and Weidner’s clocks, poorly with telomere length, with Pace of Aging and DunedinPoAm, but relatively stronger with Levine’s clock and homeostatic dysregulation, both methods that employ biomarker data [27, 28, 30, 31, 149, 367]. In a study of 11 diverse epigenetic clocks including biological age measured by KDM, Belsky concluded that different types of clocks are capturing different features of the aging process [30]. Biological age as measured by KDM was found to be superior to chronological age in predicting risk of diabetes, stroke and cancer, while chronological age was better suited to predicting risk of dementia and Alzheimer’s disease, and both methods were deemed equally effective at predicting risk of coronary heart disease and all-cause mortality [430]. Work with the ACCORD trial detected a twelve year older biological age as calculated by KDM in diabetics as compared to healthy controls; prediabetics were 2.69 years older than healthy non-diabetics [14]. Similarly, work with the CARDIA study population detected higher odds of both cardiovascular disease and stroke associated with elevated biological age [117]. Comparing control individuals versus those who experienced either a major coronary event, ischemic stroke, or intracerebral hemorrhage found that all three adverse medical events were associated with elevated biological age in a Chinese cohort [66]. KDM-based biological age has also been associated with multimorbidity and all cause mortality as well as specifically mortality due to cancer and diabetes [14, 66, 77, 94, 117, 139, 361]. However, a study of hospitalized COVID-19 patients found no association between biological age and mortality due to COVID-19 infection [387]. KDM-measured biological age has been found to differ between men and women, with different formulas being calculated for each sex in several studies [175, 467]. There may be different racial components to KDM-measured biological age as well: in a 2014 study using 10 biomarkers, it was found that African American participants in the NHANES study showed a higher biological age than did their white counterparts, leading to an overall 46% increase in risk of mortality, 40% more likely to die due to cardiovascular disease, and 51% more likely to suffer fatal cancer [220]. A pair of studies published in 2022 support this finding, observing that Black study participants were biologically older than white participants, and in the CARDIA study, an increased biological age in Black males led to a significantly increased risk of cardivascular disease as compared to white males [117, 139]. A 2021 study using 12 biomarkers found that the KDM calculation on those features were predictive of mortality [238]. However, a KDM model trained on NHANES data was found to be predictive of mortality in a Taiwanese study population, suggesting that while different racial populations may exhibit different rates of KDM aging, there may still be common trends in biomarkers across racial groups [129].

Further, KDM-based biological aging rates are not static: a 2018 paper describes a decline in KDM-measured biological age acceleration in later generations, largely attributable to modifiable health-related behaviours including smoking cessation [222].

Physical functioning is associated with KDM-calculated biological age, with declines in physical ability associated with higher biological age [25, 30, 149]. Study in the HRS population detected an association between prevalent and incident limitations in activities of daily life with elevated biological age, a finding that was significantly exacerbated in Black versus white study participants [139]. A 2022 study reported that worse physical and emotional health was associated with elevated biological age, although curiously they also reported finding grip strength inversely associated with KDM-calculated biological age [150]. Similarly, declining cognitive function is associated with higher biological age [25, 30, 77, 129, 149]. Short-delay free recall, self-reported health and self-reported disability, subjective aging parameters such as facial aging, and increased comorbidities, hospitalization, and difficulty with basic and instrumental activities of daily life also associate with KDM-based biological age [30, 149, 467].

Mental health impacts including perceived stress, experienced discrimination and depression also are linked to elevated biological age [118, 150]. In a pair of studies using partially overlapping sets of inflammation-related biomarkers, Lin and colleagues studied whole blood gene expression changes associated with changes in biological age as determined by those biomarkers [231, 232]. In studies of the FHS Offspring cohort, they found 448 genes whose expression level changed in response to biological age change as measured by 7 inflammation-associated biomarkers, and 481 genes whose level changed in association with biological age as measured by 6 biomarkers in a second study two years later. Urinary markers of oxidative stress were used to develop a KDM-based biological clock, indicating that urinary metabolites are useful as a data source for biological aging [285]. Biological age as calculated by KDM was lower in calorie-restricted diet adherents versus those with no restriction [29]. A study of macronutrient consumption found that above average consumption of carbohydrates in conjunction with typical consumption of proteins and lipids exhibited the lowest level of biological aging among dietary compositions [358]. Inflammatory dietary potential is positively associated with biological age acceleration in a study of NHANES participants [446]. In a study of obese older adults, it was found that either diet alone or diet in conjunction with exercise reduced biological age acceleration more than did exercise alone, or control [158]. Related to diet, hydration was found to associate with biological aging: higher serum levels of sodium, a proxy for hydration habits, resulted in a worsening KDM-calculated biological age [94]. In a study of pre- and postmenopausal women, it was found that parity was associated with different rates of biological aging as measured by KDM, but only in postmenopausal women [367]. A concave relationship between parity and biological aging rate was observed, with the lowest rate of biological aging observed for women who had 3 or 4 live births.

Lastly, a pair of studies assessing the relationship of socioeconomic and lifestyle features with KDM-calculated age were published in 2020, both making use of the Singapore Longitudinal Aging Study population. Both studies observed the age decelerating effects of increased education, although other features were observed differently in each study [290, 467]. In the study undertaken by Ng and colleagues, they observed an effect from nearly all of the variables considered, but noted that accelerated biological aging was especially associated with being single, widowed or divorced, with living alone, smoking, elevated alcohol consumption, being at high nutritional risk or exhibiting poor nutrition [290]. They found some gender differences, with housing type and unintended weight loss being a common feature for both men and women, but marital status being more significant for men and diet being more significant for women. In the study undertaken by Zhong and colleagues, they also found that marital status was significant for men but not women, but also observed calculated scores for health activity, social activity, productivity activity, recreational activity and cognitive stimulation were significantly associated with biological age [467]. Sleeping patterns including snoring and insomnia are associated with elevated biological age, while the same study found that conversely, an early chronotype and normal (neither reduced nor excessive) sleep duration were linked to improved biological aging [126]. Environmental parameters including neighbourhood resource availability and air pollution associate with KDM-calculated biological age [118, 306]. A 2023 study observed that lower socioeconomic status exacerbated mortality risk associated with elevated biological age [361].

#### Novel and Specific Clocks

In addition to the aforementioned clocks, there are a wide variety of lesser used epigenetic clocks as well as recently developed clocks, often for particular purposes. Here we offer a brief overview of these clocks. Five studies reviewed here made use of the seven CpG Vidal-Bralo clock, finding its correlation with chronological age and telomere length, as well as its increased precision in men relative to women [15, 156, 410, 413]. A pair of studies making use of the Vidal-Bralo clock in the BASE and BASE II populations found that while the seven CpG Vidal-Bralo did not correlate with morbidity, lung capacity, or subjective health, a modified eight CpG version of Vidal-Bralo did exhibit a weak association with cardiovascular health [97, 217]. It was found that men, but not women, with obstructive coronary artery disease exhibited an elevated biological age by the Vidal-Bralo clock [16]. A 2020 study comparing the efficacy of 14 epigenetic clocks included Vidal-Bralo as well as the chronological age-trained Lin, Garagnani and Bocklandt clocks, the mortality-trained Zhang clock, and the mitotic division-trained MiAge and Yang clocks, and found that mortality-trained clocks showed accelerated aging in schizophrenics, while mitotic clocks showed decelerated aging, and chronologic age-trained clocks found deceleration among male schizophrenics taking clozapine or a combination of clozapine and an additional antipsychotic [156]. The Lin clock was found to be significant for women with elevated depressive symptoms, but a second study found no link with that clock to socioeconomic status [37, 230, 354]. The Yang EpiTOC clock was similarly found significant for depressive symptoms but was also found to be significant in the HRS but not MESA populations for association with socioeconomic status [37, 354]. The Zhang mortality-trained clock was additionally significant for depressive symptoms [37]. The EpiTOC and MiAge clocks were examined in a study of obese and overweight breast cancer survivors undergoing weight loss, weight loss therapy supplemented with metformin, or metformin alone, but no significant difference in biological aging by those clocks was detected after six months [296]. The Wu epigenetic clock was the only clock among a variety tested to detect a biological age difference in pre-eclampsia affected pregnant women as compared to normotensive controls [319]. In a study of French centenarians and supercentenarians and their offspring, four clocks of limited CpGs (all used 8 or fewer methylation sites) found that both the centenarians/supercentenarians and their offspring were younger than their chronological age by those clocks [86].

Zhang and colleagues developed a 10 CpG mortality score, that while not correlated with chronological age, did correlate with mortality and frailness [462]. The Health Aging Index (HAI) was used in a 2022 study along with KDM-calculated biological age and homeostatic dysregulation measures to identify an improvement in HAI among older obese adults who adhered to a diet and exercise combination weight loss treatment [158]. Intrinsic Capacity (IC) score, which is based on cognitive and physical performance as well as geriatric depression diagnosis associated a lower baseline IC with higher inflammatory marker levels, particularly TNFR1 and GDF15 levels [246]. The TruMe proprietary clock, which exhibits a good correlation with chronological age, was used to detect a decrease in biological age following seven months of a pharmaceutical interventionf [89]. The MARK-AGE project biomarkers, which include DNA methylation as well as serum metabolites, peptides, and N-glycans, was used to analyze a population of HIV-positive and control blood donors [88]. That study found many of the MARK-AGE markers elevated in HIV-positive participants versus controls, meaning that the HIV-positive participants were significantly biologically older, particularly male participants. The Aging 3.0 clock has also been used to study the HIV positive population; a 2023 study reports elevated age among cannabis but not tobacco smokers, as well as increased biological age in diabetics as compared to non-diabetics [254, 313]. BrainAge, which associates with chronological age positively, was also found to associate positively with depressive episodes, their duration, and age of onset [424].

Several studies analyzed for this review made use of a metabolic-based epigenetic clock, using markers of metabolic syndrome to calculate biological age. In a pair of studies published in 2018, Han and colleagues studied metabolic syndrome status-related biological age based on waist circumference, blood pressure, fasting blood glucose, serum triglycerides, HDL cholesterol, total cholesterol, and chronological age [145, 146]. They found increased biological age by this metric associated with male sex, increased BMI, chronic disease diagnosis including hypertension and diabetes, less than six hours of sleep nightly, lower educational attainment, marital issues, unemployment, low income, lack of exercise, being an ex-smoker and low consumption of alcohol, while decreased biological age was associated with higher usage of nutrition data and higher socioeconomic status. A 2018 study based on 18 physical parameters and biomarkers developed a biological aging clock that indicated increased risk of mortality, hypertension, diabetes, stroke and cancer with an increased biological age [180]. In a study of 263,000 participants, metabolic syndrome features were analyzed using PCA, revealing different aging formulas for men and women [179]. Waist circumference, mean blood pressure, fasting blood sugar, serum triglycerides and HDL levels were all strongly associated with metabolic syndrome-based biological age, although to different degrees in men and women. An additional study established a metabolic syndrome-adjacent ‘body shape clock’, based on waist to hip ratio, hip circumference, lean body mass percentage, weight, and height, calculating different biological age formulations for men and women [13]. Waist to hip ratio was found to be most strongly associated with chronological age among the markers considered, with hip circumference and height also showing strong association. Plasma proteomics have also been used to develop metrics of biological aging: in a study of postmenopausal women, SASP-associated proteins were found to be indicative of the biological aging process [366]. Using a stacking model and 19 biomarkers, Yang and colleagues developed a novel clock with a MAE of 4.4 years in the test data set, which indicated accelerating biological age with increasing levels of comorbidity [448]. Similarly, a clock using the XGBoost algorithm and only five biomarker features found association with both all cause and cause specific mortality [426]. Using a limited number of CpG sites and measurements of signal joint T cell receptor arrangement excision circles (sjTRECs), Paparazzo *et al.* were able to produce a blood based novel clock with a mean absolute error (MAE) of 4.43 years [307]. In a study of Polish men, a body shape-based biological age was calculated based on ponderal index, waist circumference, chest index, pelvis-acromial index and total body water [136]. That study found that biologically older participants showed poorer performance on a battery of physical tests than did the biologically equal or younger participants. A separate study that considered female residents of Tiraspol used morphofunctional parameters including physical parameters and grip strength to divide the study into ‘slow aging’ and ‘fast aging’ subgroups found that most of their study participants classified as ‘slow aging’, and that their especially long-lived study participants were more likely to exhibit a lower BMI, shorter height, lower grip strength, lower skeletal muscle mass and fat mass, and higher systolic blood pressure [289]. In a study of young women, a clock based on physical parameters and body shape revealed delayed biological aging associated with elevated weekly physical activity [195]. A ’physical fitness age’ was found to be a better predictor of sarcopenia than was chronological age in a Japanese study, and a deficit in physical fitness age was found to be associated with a history of diabetes, obesity, and depression [400].

In 2022, a study was published comparing a novel measure of physiological age (based on frailty, comorbidities, and limitations in activities of daily life) across different nations, revealing a worse physiological age among Italians, Israelis and Americans, while Swiss, Dutch, Greek, Swedish and Danish citizens exhibited an improved physiological age [326]. This study also found an association with their measure of biological age and education, income, and marital status. Using a multidimensional measure of disease status, risk factors, socioeconomic and functional status along with blood based biomarkers, a 2023 study developed a clock that was more accurate than chronological age at predicting mortality [347]. Studies of facial photograph-based perceived age was found to be linked with body adiposity but not glycemic markers in a paper suggesting perceived age as a proxy for biological age [456]. Other image-based studies have focussed on skin, specifically the hands or the skin surrounding the eye; both found a positive association for their skin-based biological age and chronological age, with skin aging possibly serving as a proxy for systemic oxidative stress [38, 464].

A number of studies explored analysis of clinical biomarkers with methods such as PCA or MLR, often in comparison with KDM. Several studies compared the three approaches, typically finding KDM the most accurate method [173, 219, 467]. A 2013 study found that KDM outperformed MLR, which in turn outperformed PCA, while a 2020 study found that while KDM was best at predicting prevalent frailty, PCA and MLR were competitive in terms of predicting frailty incidence and mortality [219, 467]. A 2017 study found that while KDM-calculated biological age best correlated with chronological age, and that all three metrics showed increased biological age associated with increased glucose tolerance, while PCA and KDM showed elevated biological age for those diagnosed with diabetes [173]. A 2023 study of 9 biomarkers tested the performance of PCA, MLR and KDM to predict biological age, and found that PCA and MLR outperformed KDM-based calculations in terms of correlation with chronological age [229]. In a series of three published studies, Zhang and colleagues use PCA to select different sets of biomarkers to estimate biological age in a Han Chinese population [458, 459, 461]. Using 18 biomarkers and PCA, a 2018 study describes a biological age metric developed with a Korean study population, where they observed increased mortality and increased risk of hypertension, diabetes, heart disease, stroke, and cancer with increased biological age [180].

In a pair of studies, Pyrkov and colleagues explore the possibility of locomotor activity-based metrics of biological aging; in their work, they determined that a PCA-based locomotor activity clock outperformed other algorithms as well as a convolutional neural network trained on the same data [320, 321]. In a study comparing the KDM-based analysis of markers described in Levine’s 2013 paper, as well as a set of phenotypic markers and the TAME assays, all three measurements were found to be associated with increased difficulty in activities of daily life, multimorbidity, cognitive dysfunction and mortality in the HRS population [78, 219]. Methylation of 10 mortality-associated CpG sites were used to calculate a ‘Mortality Risk Score’, which correlated with both Horvath’s pan-tissue and PhenoAge epigenetic clocks, as well as inversely with telomere length [125]. Measured on a ten point scale, increases in the Mortality Risk Score was associated with increased all-cause mortality, fatal cardiovascular disease, and fatal cancer. Using the NHANES study participants and a set of 17 selected biomarkers, Rahman and Adjeroh describe an exploration of MLR, KDM and Deep Neural Networks (DNN) in conjunction with centroid- or medoid-based (mean versus median) age neighbourhood construction, finding that the DNN performed best, with medoid-based estimation performing best for mortality modeling [324]. A study developed using the Canadian Longitudinal Study on Aging made use of the KDM calculation to develop six interrelated measures of biological aging, corresponding to different physiological systems, which showed different rates of aging in men and women, and different associations with mortality, multimorbidity, and hospitalization [409]. Research has also explored the predictive capacity of physical ability on biological age. The functional aging index which is composed of gait speed, subjective sensory ability, muscle strength and lung function, was found to significantly increase risk of entry into care and mortality, with one study showing an associated mortality risk of 27% per standard deviation of functional aging index-measured biological age [113, 228]. That study also considered the frailty index and cognitive function, finding that while functional aging index correlated with frailty index, it negatively correlated with cognitive function [228]. The study also associated 32% and 15% increased mortality risk per standard deviation of biological age measured by frailty index and cognitive function respectively. An additional study found that frailty index was a better predictor of mortality than Horvath’s pan-tissue clock [188]. Using 29 indices of cardiovascular function and testing four different machine learning approaches, Schumann and colleagues found that the Gaussian process regression model outperformed other approaches in terms of estimated biological age and chronological age correlation, and found that cardiovascular changes were associated with obesity in accelerated biological age by that measure [356]. One paper described the use of fBioAge, which is a composite of grip strength, FEV1, vision and hearing [384]. The study found that the ‘slow aging’ group tended to be older, more likely male, had more schooling, better literacy and fluency, better recall, processing speed, and semantic knowledge. Immunity-based an immune cell proportion clocks have also been developed in recent years, with three papers reporting clocks based on immune system features. The 2021 ImmunoAge clock found an association between anxiety and elevated ImmunoAge in women, while healthy centennarians exhibited a lower ImmunoAge [262]. That paper also reported a one month nutritional intervention resulted in an improved ImmunoAge. Using an immunity-based clock in a study of men with clinically stable COPD and controls, it was found that COPD patients have an elevated immune based age due to a combination of immunosenescence, pro-inflammatory state and oxidative stress [265]. A 2023 paper used partial least squares regression on immune cell proportions and found delayed age-related changes in those cell type proportions in supercentennarians; they also observed an association with lower inflammation in those study participants [468].

Lastly, we address the novel, often purpose-focused clocks we examined for this review. A novel clock based on clinical parameters associated with cardiovascular function has been developed using linear regression techniques, finding sex-specific associations with different clinical features and cardiovascular health [111]. Two clocks dedicated to assessing biological age in brain tissue were developed, with very different technical approaches [368, 378]. A 2015 paper describes the development of a ‘healthy aging score’ based on RNA levels of 150 genes, in which a better healthy aging score was found to be associated with better renal function in a 12 year follow-up [378]. Post-mortem brain samples from Alzheimer’s patients exhibited a lower healthy aging score, as did samples from individuals with mild cognitive impairment. Following the observation that many epigenetic clocks systematically underestimate the age of brain cortex samples, Shireby and colleagues developed a novel clock that more accurately assesses biological age in cortex samples [368]. This clock employed elastic net regression to select CpG methylation sites significantly associated with chronological age. Other novel clocks have been developed using elastic net regression, including a skin-specific, a breast-specific, a skeletal muscle clock, and a metabolomic age clock [41, 60, 339, 415]. The first three use CpG methylation while the fourth makes use of serum and urine biomarkers as assessed by liquid chromatography-mass spectrometry and nuclear magnetic resonance spectroscopy; that clock was trained on a population of men, and found that their developed metabolomic age correlated with chronological age, and that elevated metabolomic age was associated with being overweight or obese, habitual heavy drinking, presence of diabetes, depressive symptoms/depression, anxiety and post-traumatic stress disorder [339]. The skin-based clock was developed using methylation signatures from 2266 CpG sites, and was found to work well on both biopsied samples and tissue culture [41]. The breast-based clock relies on data from 286 CpG sites, and when tested on normal breast tissue highly correlates with chronological age [60]. When tested on breast tumour samples, those samples show biological aging acceleration regardless of HER2 and hormone receptor genotype, while for triple-negative breast tumours no apparent acceleration was observed. The skeletal muscle-derived clock relies on the contributions of 1975 CpG sites, and outperforms Horvath’s pan-tissue when it comes to estimating biological age from that tissue source [415]. The GlycanAge clock, based on measurements of IgG glycan structures, found that among 24 glycan structures measured, 21 of them were significantly associated with changes in chronological age, from which the authors developed their clock [201]. They found that galactosylation was the most significant age-related glycosylation change, and that men and women exhibit different patterns of change in glycosylation.

Medical imaging has also been explored for novel clock development. The chest radiograph based ’CXR-Age’ clock is associated with baseline cardiovascular disease risk factors including male sex, diabetes or hypertension diagnoses, obesity, and smoking status [322]. Two retinal imaging based clocks have been developed: 2022’s ’RetiAge’ clock associated with all cause mortality, cardiovascular and cancer mortality, while a novel clock based on retinal microvasculature published in the same year finds that this non-invasive data source may reveal important information on biological aging and cardiovascular disease risk [292, 369].

Finally, we address deep learning-based clocks, trained on a variety of input data including blood biomarkers, physical behaviour, and facial appearance [254, 255, 323, 324, 442]. A pair of studies demonstrated that deep learning-based methodologies are highly competitive with other machine learning approaches such as regression, finding that with a large training set, deep neural networks could achieve comparable correlations and mean absolute errors [254, 255].

Using deep neural networks (DNNs) on three genetically distinct training populations observed that their trained models performed best on the populations they were trained on, with decreased performance on other populations, implying that there were population-specific significances associated with the blood-based biomarkers used as features [254]. A deep neural network trained on non-smokers measured accelerated aging in a smoking test population, again based on blood derived biomarkers [255]. A DNN clock based on 36 markers with a MAE of 6.0 years demonstrated improvements in measured biological age following adherence to either the Mediterranean or DASH diets [106]. A similarly accurate DNN-based clock developed using the Moli-Sani population found that elevated biological age was associated with all cause mortality, hospitalization, cardiovascular disease, ischemic heart disease and cerebrovascular disease [134]. A distinct clock developed from the same study population determined that elevated dietary inflammatory index and dietary inflammation score resulted in in increased biological age [261]. An additional DNN-based clock, based on 16 biomarkers and exhibiting a MAE of 5.68 years, revealed an association of stroke history, liver disease and lung conditions, but no association for heart disease or cancer history, with elevated biological age [124]. This clock also revealed that smoking, marital status, and sleep quality were key factors in influencing biological age. Using a combination of chronological age and locomotor activity collected from the NHANES data set, applied to a convolutional long short-term memory (ConvLSTM) neural network, this network could predict biological age more accurately than could one dimensional convolutional neural networks (CNNs) or deep neural networks [323]. A study employing a back propagation artificial neural network (BP-ANN) to develop a biological aging clock achieved a correlation of 0.68 with chronological age, and found that a diet rich in both polyphenols and proboiotics showed a decrease in biological age acceleration [279].

Lastly, the CNNPerceivedAge CNN can not only determine chronological and perceived age from a photo, and could also associate health parameters such as BMI, blood pressure, and clinical blood biomarkers with that photo [442].

## Discussion

While the past decade has seen substantial increases in both the breadth and depth of research into the question of biological aging, including the development of many novel approaches to measurement, there are both outstanding and newly generated questions yet to be addressed. We break these questions into five categories: questions regarding measurement, questions regarding biological aging itself, questions of group differences including ethnicity- and sex-based differences, methylation measurement-specific and biomarker-based measurement questions.

### Measurement Questions

It has been observed by some of the papers here reviewed that differing measures of biological age, such as telomere length, mitochondrial DNA copy number, methylation, or blood-based biomarkers do not always concur. This leads one to question whether there is a direct relationship between these different measurements and if so, what the nature of that relationship might be. What biological processes might underlie these very different molecular features? Indirectly related to this question is the characterization of different markers of biological age as responsive to aging, while which are generative of the aging phenotype. Differentiating between these two feature types would greatly facilitate both intervention and diagnostic efforts.

For different environmental or health conditions, understanding the influence of those conditions on the expression of biological aging remains to be more fully characterized. It is unclear whether these conditions are genuinely influencing biological aging rates, or if they are influencing the measurement of biological aging. This question relates to the responsive versus generative dichotomy of biological age markers, as those conditions that change the expression or status of aging generative markers would authentically modify biological aging trajectories, while those that influence responsive markers could be reflective only of changes to the measurement of biological age.

A more specific question regarding measurement of biological age focuses on the use of MRI-based imaging to assess changes in the brain and peripheral nervous system. Do these approaches overcome the difficulty of directly sampling those tissues, or are volumetric and morphological changes not sufficiently specific to the process of biological aging? Which changes are informative, while which are less specific?

### Methylation-based Measurement Questions

Many measurements of biological age use CpG methylation and the focus on this molecular feature has led to the generation of a number of possible future research questions. Pre-eminently among those questions is: why is there a markedly limited overlap in the CpG sites employed by different methylation clocks? Is this limited overlap due to different tissue and cell types used for clock development, meaning that there are different markers of biological aging in different tissue types, or is there another explanation? Many, but not all of these methylation clocks show slower aging in female subjects, despite the relative acceleration of observed aging in female-specific tissues such as mature breast. Why does breast tissue typically display accelerated aging relative to the whole individual and, related, why would other (typically blood-based) clocks show decelerated aging in females relative to males? Further, if different tissue-based methylation clocks correctly measure different rates of aging in their respective tissues, does this imply that a generalized, blood-based methylation clock is inherently limited in precision for a heterogeneously aging organism? If so, does this imply that multi-tissue biomarkers are a more valid approach to the measurement of biological aging? Does this also imply that imprinted genes, that is, those genes whose alleles exhibit differential methylation based on the parent of origin, are also directly implicated in the process of biological aging under both normal and pathological conditions?

In addition to sex-based differences in methylation-based age measurement, there has also been observed differential CpG methylation in different ethnic groups, with most of the relevant work comparing African American and Caucasian individuals. Does it also follow that all ethnic groups have a group-specific set of CpG sites that are methylated differentially during aging? Conversely, is there a set of ethnicity-independent CpG sites that reflect the status of biological aging for all humans?

### Biomarker-based Measurement Questions

Some of the biological aging clocks reviewed here make use of blood-based biomarkers including specific serum protein levels and metabolites. Following from our previous questions regarding sex- and ethnicity-specific methylation differences, why do some biomarkers exhibit significant sex-based differences in relation to biological aging? Do these reflect a difference in biological aging rates between the sexes, or is it independent of the rate of biological aging? A number of biomarker-based studies are done in Caucasian and Asian populations but with one exception to be discussed shortly, we have not found any systematic research comparing biomarkers between ethnicities in the time period reviewed here. Are blood-based biomarkers ethnicity-dependent as well, or are they less dependent on ethnicity than CpG methylation appears to be?

### Other Measurement Questions

One paper compared biomarker-based ethnic differences between three distinct groups [254]. This paper employed a deep learning-based approach to assessing biological age in large populations, including distinct models trained on each of the groups considered, and then tested on non-training data. An additional paper used deep learning trained on facial photographs to assess biological aging [442]. While neural network and deep learning approaches are undoubtedly powerful, are they at risk of learning features other than biological aging, for example, underlying ethnic differences? Beyond sex- and ethnicity-based differences, such as socioeconomic status or cultural differences in diet and lifestyle, there are further potential confounding factors that may stymie the development of a so-called universal biological aging clock. Given these factors, is a pan-ethnicity biological clock genuinely possible, or will different sub-groups respond more accurately to bespoke measurements?

Within larger ethnic groups, different patterns of biological aging are also observed. In the work of von Kanel and colleagues, it was consistently found that Black South Africans displayed shorter telomeres than their Caucasian colleagues, contrary to findings in American populations [417, 418]. What underlies these differences in observations? Do other markers of biological aging in South Africans diverge from patterns observed in Americans? Investigating these differences may help to tease out the influences of non-biological factors on patterns of biological aging.

### Biological Age Variables

It has long been speculated that diet and lifestyle, among other environmental and modifiable features, contribute to differing rates of biological aging. Is diet a primary contributor to biological aging, or rather, is it an indicator of other socioeconomic factors that themselves directly contribute to differences in biological aging? Similarly, physical activity has been found to modulate biological aging, but the underlying relationship between physical activity and biological age remains to be characterized.

Some of the papers reviewed here explore biological aging changes in relation to pregnancy and gestation, while other work has considered the contribution of early life events to later biological aging status. What contributions do maternal health make to later biological aging? How do early life events influence biological aging: do they alter biological aging rates at the time of the event, or is it a delayed effect? In either case, is there an acute feature, ideally a molecular or epigenetic feature, that can assess the biological aging contributions of a given early life event?

While telomere length has been explored in a number of model organisms including non-vertebrate organisms, not all measures of biological aging have been characterized in model organisms. Are biological aging markers such as gene expression and methylation status a human-specific, primate-specific, mammalian-specific, or general animal feature of biological aging? The answer to this expansive question will greatly influence what animal models are most relevant to the study of aging in human subjects.

## Conclusion

This review has taken a careful examination of a twelve and a half year window of work pertaining to the study of biological aging, focusing on methodological developments and molecular features. While a great breadth and depth of work has been undertaken, numerous questions remain to be answered, while new questions and testable hypotheses have been additionally generated. A number of these outstanding questions pertain to the universality of a hypothetical biological clock, or whether such a clock is not strictly possible. Our current body of knowledge and methodology sets the stage for exciting possibilities in further understanding of the process of biological aging, along with opportunities for early surveillance and intervention.

